# 20E-dependent tyrosine phosphorylation of phospholipase C gamma underpins egg development in the malaria vector *Anopheles gambiae*

**DOI:** 10.1101/2022.09.01.506175

**Authors:** Simona Ferracchiato, Flaminia Catteruccia, Matthew J. Peirce

**Affiliations:** Dipartimento di Medicina e Chirurgia, Università degli Studi di Perugia, Perugia 06132, Italy; Department of Immunology and Infectious Diseases, Harvard T.H. Chan School of Public Health, Boston, MA 02115, USA; Howard Hughes Medical Institute, Boston MA, USA

**Author notes:** Co-corresponding authors (MP), (FC).

## Abstract

The efficient development of eggs following a blood meal is central to the vector capacity of *Anopheles gambiae* females. The ecdysteroid hormone 20-hydroxyecdysone (20E) plays a pivotal role in this process, yet the signaling mechanisms by which 20E exerts its effects remain incompletely understood. Here we show blood feeding is associated with increased tyrosine phosphorylation of an array of proteins coincident with the increase in titers of 20E that follows a blood meal. Injection of genistein, a tyrosine kinase inhibitor, reduces both the appearance of phosphotyrosine-containing proteins and fecundity, linking tyrosine phosphorylation to egg development. We identify one of the proteins phosphorylated after a blood meal as phospholipase C gamma (PLCγ) and show that its blood feeding-induced phosphorylation is dependent on endogenously synthesized 20E via the ecdysone receptor (EcR). Interruption of Src-like tyrosine kinase signaling inhibits phosphorylation of PLCγ as well as egg development after blood feeding, implicating Src-like kinases in the phosphorylation of PLCγ. Taken together, these data suggest that the effects of 20E on *An. gambiae* egg development are at least partially mediated through a tyrosine phosphorylation-dependent signaling cascade involving Src family tyrosine kinases and PLCγ.

## INTRODUCTION

*Anopheles gambiae* females are one of the most important sub-Saharan vectors for *Plasmodium falciparum* parasites which cause malaria, a disease which endangers half the world’s population and resulted in more than 600,000 deaths in 2020 (1). The capacity of *An. gambiae* to transmit *Plasmodium* parasites is underpinned by its blood feeding behavior. Blood feeding provides the nutritional resources the mosquito needs to generate eggs (oogenesis), thus sustaining a large vector population that supports disease transmission. Moreover, the resources provided by blood feeding favor the growth and development of *P. falciparum* within the mosquito (2, 3). Specifically, recent data show that 20 hydroxyecdysone (20E), an ecdysteroid hormone induced in the mosquito female following a blood meal, besides being essential for the development of eggs, regulates the survival of parasites that are crossing the midgut (2). Indeed, pharmacological targeting of 20E function simultaneously attenuates both mosquito fecundity and parasite development (4). The coordinated regulation of oogenesis and parasite development has galvanized interest in defining the signaling events launched by blood feeding, and specifically those dependent on 20E, as possible targets to limit malaria transmission by reducing both mosquito reproduction and parasite survival (4, 5).

The most relevant and comprehensive analysis of the sequence of events leading to 20E production following a blood meal comes from the *Aedes aegypti* mosquito (6), vector for arboviral infections such as Yellow fever, Dengue fever, Chikungunya and Zika. Stretch receptors in the midgut, activated by the presence of blood, cause the synthesis of ovarian ecdysteroidogenic hormone (OEH) in the brain and its release from a specialized storage organ in the *corpora cardiaca* (7, 8). OEH acts on epithelial cells of the ovarian follicles driving the synthesis of the precursor of 20E, ecdysone (E) (9, 10). Upon secretion, E is taken up by the fat body where it is hydroxylated at the 20 position to generate the active hormone 20E, which acts through a heterodimeric steroid hormone receptor complex comprising the ecdysone receptor (EcR) and Ultraspiracle (USP)(11). This complex upregulates the expression of an array of 20E-regulated early genes including the transcription factors E74 (12), E75 (13) and Broad complex (BR-C) (14) which in turn control the vitellogenic transcriptional program described below (15).

In addition to the synthesis of 20E, blood feeding initiates the production of insulin-like peptides (ILPs) in the brain that act as growth factors for the developing oocyte through a single insulin receptor (16, 17) as well as collaborating with OEH to augment ecdysone synthesis in the ovaries (9, 18). Independently of insulin, free amino acids generated by the tryptic digestion of ingested hemoglobin are taken up by tissues and cells through receptor-mediated endocytosis, directly activating the target of rapamycin (ToR) pathway (19). As shown in *ex vivo* organ cultures of explanted *Ae. aegypti* fat body (a tissue analogous to the mammalian liver), ILPs, 20E and free amino acids act synergistically (20) to drive the production of yolk precursor proteins (YPPs) such as vitellogenin and lipid transporters such as lipophorin from the fat body (21), as well as the upregulation of the receptors necessary for their uptake in the developing oocyte (22, 23).

Important elements of the blood meal-derived signals driving egg development in *Aedes* mosquitoes appear to be common to *An. gambiae*. As discussed above, in these anophelines 20E as well as both components of its receptor, EcR and USP, play a major role in egg development (2). In addition, ILPs similar to those identified in *Aedes* are released from the anopheline brain in response to a synthetic blood meal (24), and stimulate ecdysteroid synthesis in explanted *An. gambiae* ovaries (25). However, while the major constituent signals driving egg development appear conserved between mosquito species, the molecular details underpinning this process and specifically, defining signaling events attributable to 20E, remains a work in progress.

In an effort to identify 20E-dependent signaling events after blood-feeding we hypothesized a role for phosphotyrosine-containing proteins. Mammalian steroid hormones such as estrogen and progesterone (26, 27), as well as 20E itself in some insect species (28), have been reported to signal through changes in tyrosine phosphorylation. In addition, tyrosine phosphorylation and tyrosine kinases have been implicated in 20E-regulated oogenesis in the distantly related dipteran insect, *Drosophila melanogaster* (29, 30). Here, we identify a blood-feeding-induced signaling relay in the female reproductive tract that is essential for egg development in *An. gambiae*, involving the 20E-dependent tyrosine phosphorylation of phospholipase C gamma (PLCγ) by Src family tyrosine kinases. These findings validate the concept that tyrosine kinases and their substrates may be useful targets for the design of novel agents for vector control.

## RESULTS

### Tyrosine phosphorylation after blood feeding plays a role in egg development

Since steroid hormones (27, 31) have been linked to the activation of tyrosine kinase signaling cascades, our efforts to understand the role of 20E in egg development in *An. gambiae* began by looking for evidence of changes in tyrosine phosphorylation following blood feeding. Three tissues were analyzed; reproductive tract (RT, comprising atrium, ovaries and spermatheca), head, and rest of body (RoB, comprising the remaining carcass minus the midgut to avoid contamination with phosphoproteins associated with the blood bolus). Using a monoclonal antibody, 4G10, which selectively recognizes phosphorylated tyrosine residues, we documented a robust and repeatable increase in the phosphotyrosine content of an array of proteins in the reproductive tract that was detectable at 6 hours and remained above base line for at least 48 hours post blood feeding (hPBF) (Fig. 1A, B). In contrast, blood feeding-induced changes in phosphotyrosine in the rest of body samples of the same females were weaker and less numerous while blood feeding had no apparent effect on phosphotyrosine levels in the head. In subsequent experiments we therefore focused on the reproductive tract.

**Figure 1.**
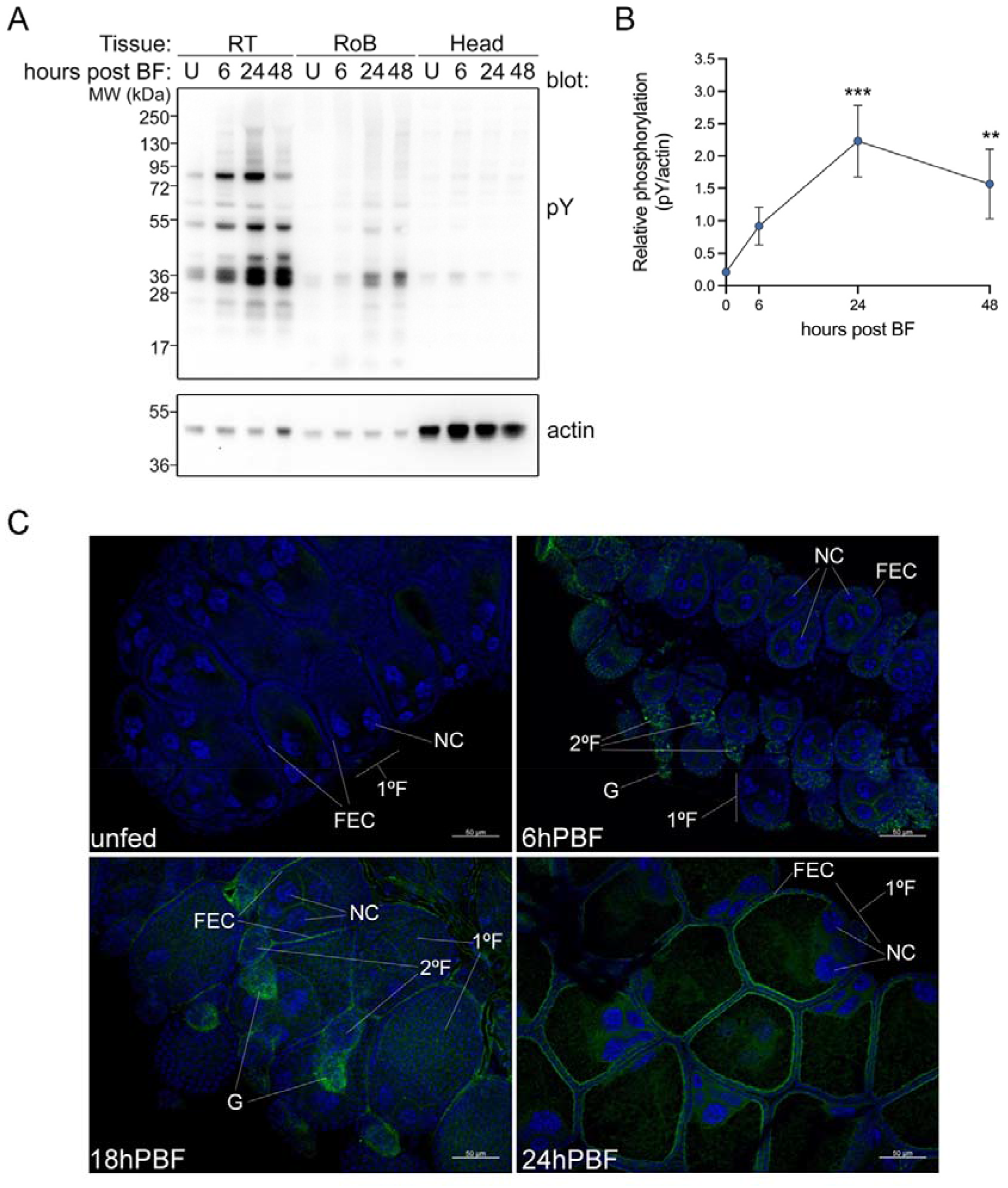
Blood feeding induces tyrosine phosphorylation in reproductive tract. A. Reproductive tracts (RT, comprising atrium, spermatheca and ovaries), heads or the rest of the body (RoB, remaining carcass minus midgut) were dissected from the same females maintained on sugar alone (unfed, U) or at the time points indicated after blood feeding. Extracts of pooled tissues (10/ time point) were analyzed by western blot for phosphotyrosine (pY), then stripped and reprobed for actin. B. Optical densities of the total pY signal in reproductive tracts were normalized against the actin signal in the same lane and expressed as ‘relative phosphorylation’. The mean ± sem of nine experiments similar to the one in A are presented in B. Changes in relative phosphorylation at each time point were compared to unfed values using a Kruskal-Wallis test with Dunn’s correction for multiple comparisons. ** denotes p<0.01, *** denotes p<0.001. Significance was ascribed to p<0.05. C. Three-day-old females were maintained on sugar solution alone (Unfed) or allowed to take a blood meal and ovaries dissected at the time points indicated post blood feeding. Phosphotyrosine (shown in green) was visualized using an anti-phosphotyrosine primary antibody and an Alexa 488-coupled secondary antibody; cellular nuclei were stained with DAPI (shown in blue). Primary follicles (1ºF), secondary follicles (2ºF) follicles and germaria (G), as well as follicular epithelial cells (FEC) and nurse cells (NC) are indicated. The images presented are from the same group of mosquitoes, and are representative of two independent experiments.

Given our specific interest in egg development, we went on to examine whether changes in tyrosine phosphorylation could be localized within developing ovarian follicles using immunofluorescence microscopy. Consistent with our western blotting data, the ovaries of unfed females were essentially devoid of phosphotyrosine whereas staining within the developing follicles was detected at 6, 18 and 24 hPBF (Fig. 1C). This signal was observed in epithelial cells of the secondary follicles, structures whose maturation to become the primary follicle of the subsequent gonotrophic cycle is initiated by a blood meal (32), as well as in the germarium, the collection of primordial germ cells and somatic follicular cells from which a new secondary follicle is generated. By 24hPBF phosphotyrosine staining was also prominent within the epithelial cells of the primary follicles.

Having established that blood feeding increased tyrosine phosphorylation within developing ovarian follicles we next performed experiments to address the role of tyrosine kinases in this increase and in the development of eggs. Thus, females were injected (2 hPBF) with the tyrosine kinase inhibitor genistein, or a vehicle control. Vehicle-injected control females behaved very much as seen in Figure 1; changes in tyrosine phosphorylation were apparent at 6 hPBF and persisted until at least 48 hPBF (Fig. 2A, lanes 1-5). Notably, in females injected with genistein, the onset of the appearance of phosphotyrosine-containing protein bands after blood feeding was delayed, and the signal was noticeably less intense (Fig. 2A, lanes 6-10). A similar reduction in the level of phosphotyrosine staining within developing follicles of genistein-injected females was also observed at 24 hPBF using immunofluorescence (Supp. Fig. 1A). Quantitative analysis of multiple images of vehicle- or genistein-injected ovaries 24 hPBF revealed a statistically significant difference in the intensity of phosphotyrosine staining in the two treatment groups (Supp. Fig. 1B).

**Figure 2.**
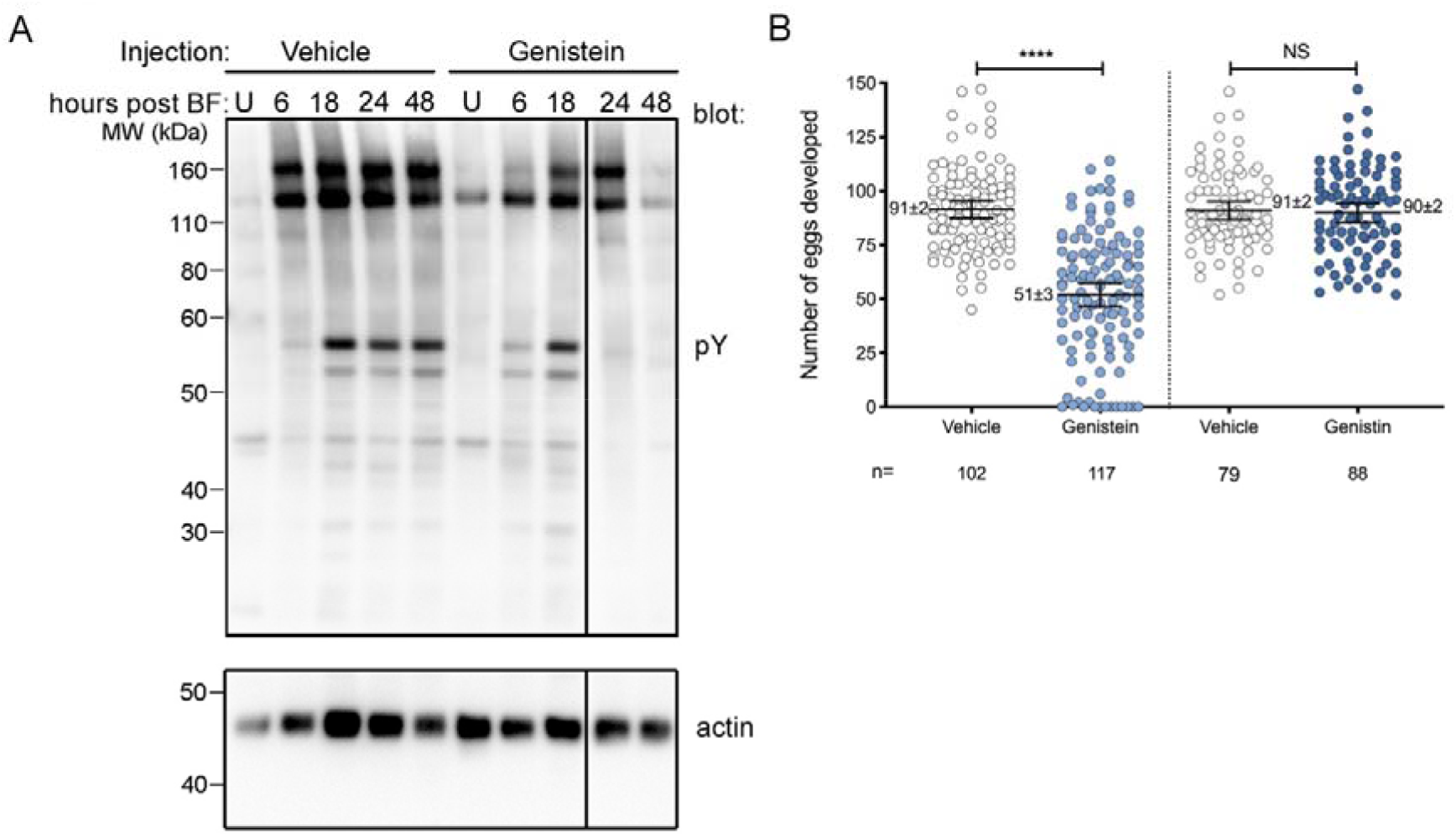
Blood feeding induces tyrosine phosphorylation in the reproductive tract that is necessary for egg development. A. Reproductive tracts (comprising atrium, spermatheca and ovaries) were dissected from females maintained on sugar alone (unfed, U) or allowed to take a blood meal then injected with the tyrosine kinase inhibitor genistein (200μM) or a vehicle control. Pooled tissues (10/ time point), dissected at the times indicated post blood feeding, were analyzed by western blot for phosphotyrosine (pY), then stripped and reprobed for actin. In the original blot the genistein-injected 24- and 48-hour time points were loaded in the opposite orientation. The line between 18 and 24 hours indicates the digital alteration of the original lane order. B. Females were injected with the tyrosine kinase inhibitor genistein (200μM) or a vehicle control (Vehicle) 2 hours after blood feeding and egg numbers were counted 48 hours later. Separately, similar experiments were performed comparing females injected with genistin, an inactive analog of genistein, or vehicle. The numbers adjacent to each group represent the mean number of eggs ± sem. Differences in the number of eggs developed in controls vs drug-treated females was compared using Mann-Whitney tests. NS denotes not significant (p>0.05), **** denotes p<0.0001. The number of mosquitoes used in the experiments (n) is shown below. Data shown represent the mean ± 95% confidence interval of 4 independent experiments with genistein and 2 with genistin.

To examine whether this tyrosine phosphorylation was required for egg development, we counted the number of eggs that developed following a blood meal in females that were injected 2 hPBF with genistein, its inactive structural analog genistin, or a vehicle control. Compared to the injection of vehicle, genistein caused a significant decrease in the number of developed eggs (Fig. 2B) whereas its inactive analog genistin had no impact. Together these data suggest a role for blood feeding-induced tyrosine phosphorylation in egg development.

### Phospholipase C gamma phosphorylation after blood feeding regulates egg development

Using a literature-based approach we next attempted to identify candidate phosphoproteins. One of the most robust changes in phosphotyrosine was observed at approximately 150kDa. We noted that in the lepidopteran insect, *Helicoverpa armigera*, 20E had been shown to cause phosphorylation and activation of phospholipase gamma (PLCγ) (28), an archetypal target of tyrosine kinase signaling which has a predicted molecular mass of 142kDa (AGAP029661). To address whether PLCγ might be a target of blood feeding-induced phosphotyrosine signaling we used an antibody specific for the tyrosine phosphorylation site (pY783 of human PLCγ1, Uniprot: <u>P19174</u>) of the enzyme that is partially conserved between mammals and *An. gambiae*. Consistent with our hypothesis, the reproductive tract of blood-fed females showed an increase in pPLCγ compared to unfed controls that was detectable between 18-48 hPBF (Fig. 3A, B).

**Figure 3.**
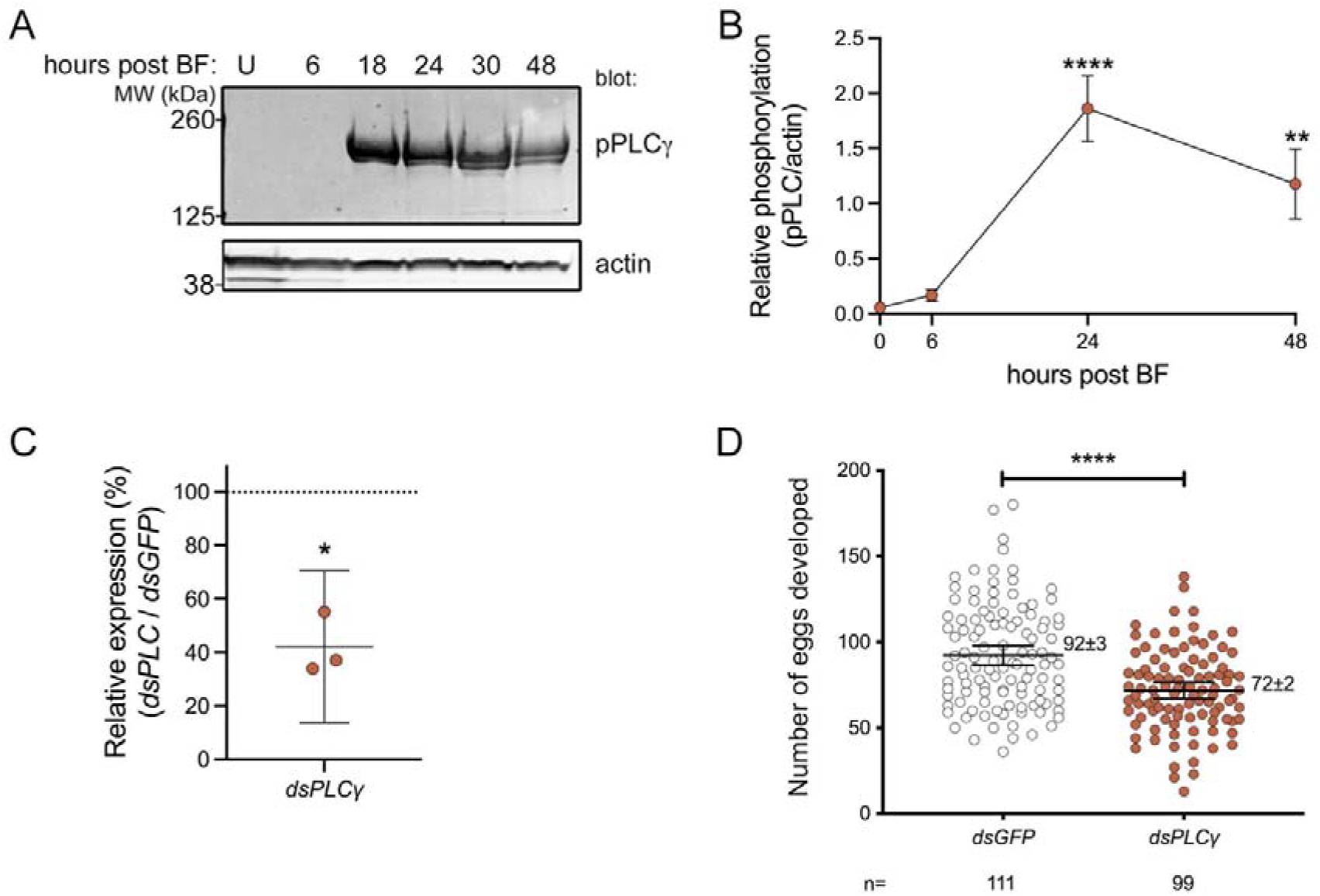
Tyrosine phosphorylation of PLCγ after blood feeding affects egg development. A. Reproductive tracts (comprising atrium, spermatheca and ovaries) were dissected from females maintained on sugar alone (unfed, U) or at the times indicated post blood feeding. Pooled tissues (10/ time point) were analyzed for phospho PLC gamma (pPLCγ) and actin. B. The optical density of the pPLCγ signal was normalized against the actin signal in the same lane and expressed as ‘relative phosphorylation’. The pooled data shown represent the mean ± sem of eight replicates of the experiment presented in A. Normality was verified using a Kolmogorov-Smirnov test and changes in relative phosphorylation at each time point were compared to unfed controls using a one-way ANOVA test with Dunnett’s correction for multiple comparisons. ** denotes p<0.01, **** denotes p<0.0001. Significance was ascribed to p<0.05. C. Expression of *PLCγ* in whole ds*PLCγ-* treated females was calculated as a percentage of the expression in ds*GFP*-treated controls in the same experiment and deviation from 100% analyzed using a one sample t test. Bars represent the mean ± 95% confidence interval; * denotes p<0.05. D. Females were injected with *dsPLCγ* or a control (ds*GFP*) then allowed to take a blood meal 72 hours later. Eggs developed were counted 48 hours post blood feeding. The numbers shown adjacent to each group represent the mean ± sem. Differences in the number of eggs developed were compared using a Mann-Whitney test. **** denotes p<0.0001. The number of mosquitoes counted (n) is shown below. Data shown represent the mean ± 95% confidence interval of 3 independent experiments.

Reasoning that this activation of PLCγ after blood feeding might signify a functional role in egg development, we assessed the impact of RNAi-mediated depletion of PLCγ on the number of eggs developed after blood feeding. Females were injected with dsRNA targeting either *PLC*γ (ds*PLC*γ) or *GFP* (ds*GFP*) as a control and then two days later allowed to take a blood meal. *PLCγ* silencing significantly reduced *PLCγ* expression (Fig. 3C) and its phosphorylation after blood feeding (Supp. Fig. 2) and impaired the number of eggs that developed compared with *dsGFP*-injected control females (Fig. 3D), suggesting that the blood feeding-induced phosphorylation of PLCγ is required for egg development.

### 20E signaling is important for PLCγ phosphorylation

We noted that the timing of the tyrosine phosphorylation of PLCγ after blood feeding matched the synthesis profile of the ecdysteroid hormone 20E, which peaks at around 24 hPBF before declining to unfed levels by 48 hPBF (33). To examine the link between the tyrosine phosphorylation of PLCγ and 20E we adopted two approaches. Firstly, we used the ecdysteroid oxidase E22O, an enzyme that catalyzes the oxidation of the critical hydroxyl moiety at carbon 22 of 20E (34), to inactivate it as reported previously (2, 35). In parallel, we used RNAi to deplete both components of the 20E receptor *EcR* and *USP*, thereby limiting 20E signaling (2, 35). Injection with E22O 2 hPBF severely inhibited the increase in pPLCγ readily detected in the control-injected females (Fig. 4A, C). Similarly, RNAi-mediated depletion of *EcR* strongly reduced the blood feeding-induced increase in pPLCγ observed in control females (Fig. 4B, D). Depletion of the 20E co-receptor *Ultraspiracle* (*USP*) had a comparable, though less robust effect (Fig. 4B), leading us to focus on EcR in subsequent experiments.

**Figure 4.**
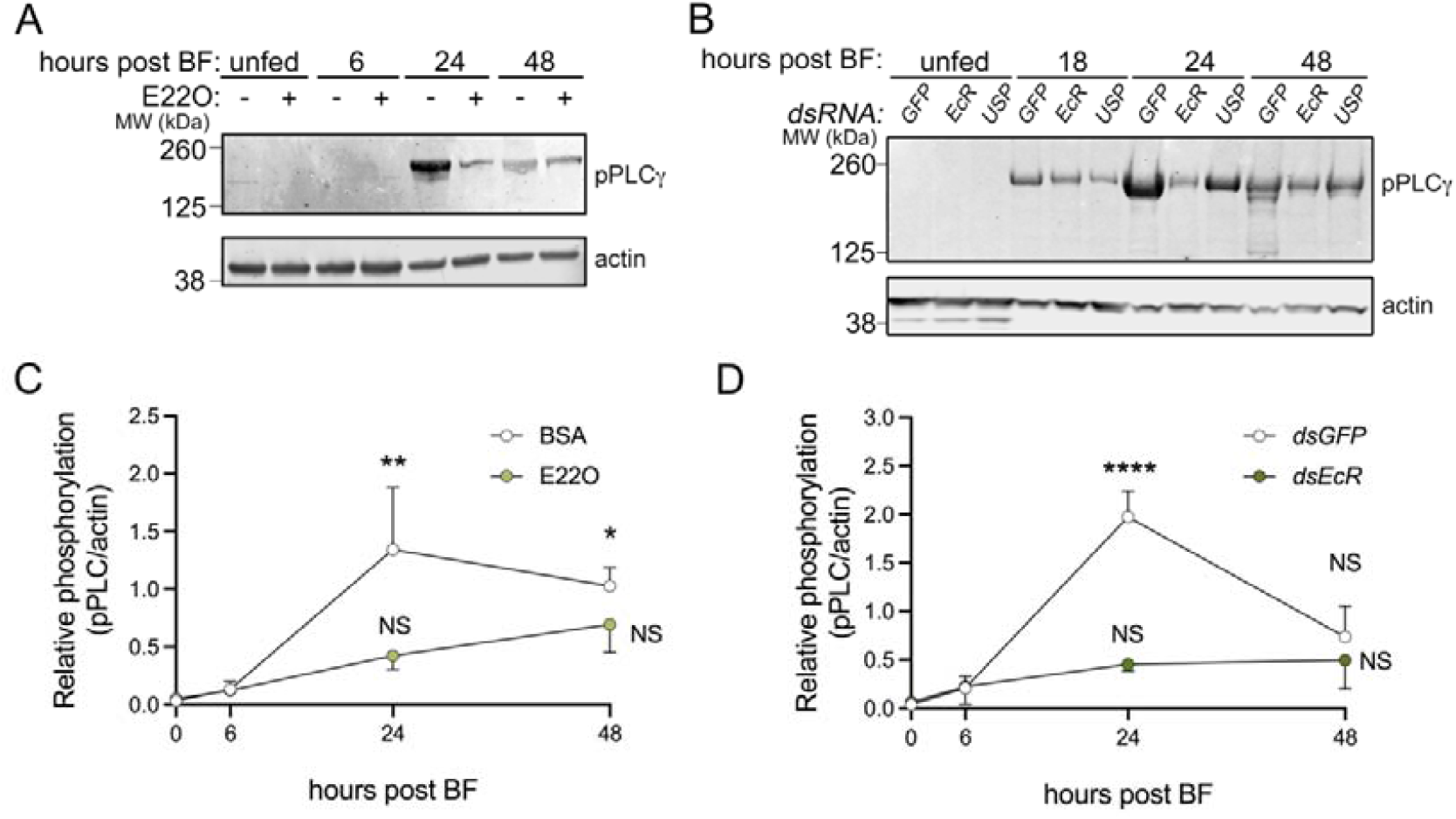
Blood feeding-induced phosphorylation of PLCγ is 20E-dependent. A, B. Reproductive tracts (comprising atrium, spermatheca and ovaries) were dissected from females maintained on sugar alone (unfed) or at the times indicated post blood feeding. Two hours after blood feeding (A) or 72 hours before blood feeding (B) females were injected respectively with E22O (5μg/μl), *dsEcR* or *dsUSP* (both 5μg/μl) or an appropriate control injection (BSA in A and *dsGFP* in B). Extracts of pooled dissected tissues (10/ time point) were analyzed for phospho PLC gamma (pPLCγ) and actin. C, D. The optical density of the pPLCγ signal was normalized against the actin signal in the same lane and expressed as ‘relative phosphorylation’. Data represent the mean relative phosphorylation values ± sem at each time point. C and D represent pooled data from four replicates of the experiments presented in A and B respectively. Differences in relative phosphorylation at each time point were compared to control unfed values using a two-way ANOVA test with Dunnett’s correction for multiple comparisons. Significance was ascribed to p<0.05. * denotes p<0.05, ** denotes p<0.01, **** denotes p<0.0001, NS denotes ‘not significant’ (p>0.05).

Given the observed effects of 20E signaling on PLCγ phosphorylation we addressed the possibility that 20E might be sufficient *per se* to induce this phosphorylation event by injecting unfed females with 20E. As depicted in Supplementary Figure 3A, injection of 20E failed to increase levels of pPLCγ in the reproductive tract, whereas blood feeding of the same mosquitoes induced a strong pPLCγ signal. Confirming the activity of the injected 20E, the heads of injected females showed a strong and time-dependent increase in the phosphorylated form of the MAP kinase c-Jun N terminal kinase (JNK) (Supp. Fig 3B), as previously reported (36). Combined, these data support a model by which 20E signaling is necessary but not sufficient for the blood feeding-induced phosphorylation of PLCγ.

### Src family tyrosine kinases are required for PLCγ phosphorylation and egg development

In many systems, PLCγ is activated directly or indirectly by tyrosine phosphorylation mediated by members of the Src family of tyrosine kinases (37, 38). To examine the possible role of this kinase family in the regulation of PLCγ after blood feeding, we used a pharmacologic kinase inhibitor selective for the Src family (SU6656) (39). Interestingly, injection of SU6656 2hPBF inhibited the tyrosine phosphorylation of PLCγ observed in vehicle-injected controls (Fig. 5A, B), whilst significantly reducing the number of developed eggs (Fig. 5C). These data implicate Src family tyrosine kinase signaling in the 20E-induced phosphorylation of PLCγ leading to egg development.

**Figure 5.**
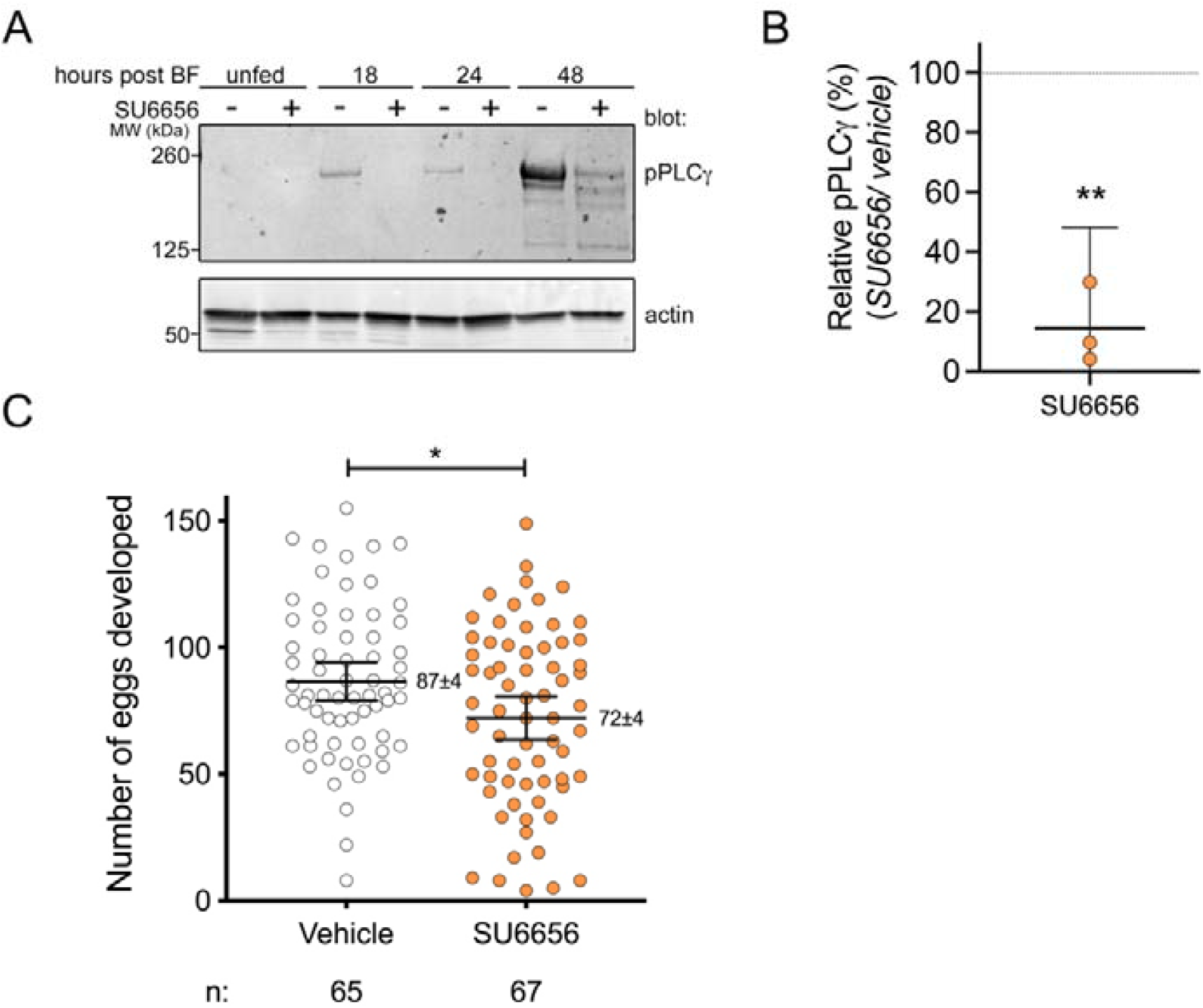
Blood feeding-induced tyrosine phosphorylation of PLCγ is partially dependent on Src-like kinases. A. Females were maintained on sugar alone (unfed) or allowed to take a blood meal then, 2 hours post-blood feeding, injected either with the Src kinase inhibitor SU6656 or a vehicle control. Females were sacrificed at the indicated times after blood feeding and reproductive tracts (10/point) were dissected and analyzed for phospho PLC gamma (pPLCγ) and actin. B. The optical density of the pPLCγ signals in panel A were normalized against the actin signal in the same lane and expressed as ‘relative phosphorylation’. The relative phosphorylation signal in SU6656-injected females was calculated as a percentage of vehicle-injected controls at 24 hPBF. Data shown represent the mean ± 95% confidence interval of three independent replicates. The difference in the pPLCγ signal between control and treatment groups was compared using a one sample t-test, all groups having passed a Shapiro-Wilk normality test. C. The number of eggs developed were counted 48 hours after blood feeding. The number of eggs ± sem is presented alongside each group and the number of mosquitoes counted (n) is shown beneath each group. The difference in egg numbers between test and control groups was determined using an unpaired t test. Data shown represent the mean ± 95% confidence interval of three independent experiments. ** denotes p<0.01, * denotes p<0.05.

## DISCUSSION

In *An. gambiae* females, the 20E-dependent signaling events that follow a blood meal regulate the development of eggs as well as of *Plasmodium* parasites (2, 3, 40). The signaling molecules involved may therefore represent novel, druggable targets for vector control. Here we characterize a signaling cascade linking blood feeding in this species to the activation of the classical tyrosine kinase substrate PLCγ, that is 20E-dependent, involves Src family kinases and is necessary for the optimal development of eggs.

We first show using both western blotting and immunofluorescence that blood feeding strongly increases tyrosine phosphorylation in the reproductive tract but not in the head or the rest of the body. The fact that genistein, a broad-spectrum inhibitor of tyrosine kinases, partially inhibits both tyrosine phosphorylation and egg development, while its inactive analog genistin does not, clearly supports the idea that the changes in tyrosine phosphorylation observed after blood feeding are important for oogenesis. To our knowledge, this is the first report of an organ-specific signature of blood feeding-induced tyrosine phosphorylation in this species despite the fact that blood feeding is known to induce the production of insulin-like peptides as well as OEH (6), both of which act through receptor tyrosine kinases (9, 41). Indeed, a possible role for these growth factors is consistent with the prominent phosphotyrosine staining we observed in follicular epithelial cells (Fig. 1C, Supp. Fig. 1A), the same cells at which insulin (42) and OEH (41) act to induce ecdysone production. Further work is necessary to confirm this possible link.

In the specific case of PLCγ the observed increase in tyrosine phosphorylation was dependent on 20E; disruption of 20E signaling, either by injection of the 20E-inactivating enzyme E22O or by silencing of the *EcR* receptor (2, 35), inhibited blood feeding-induced tyrosine phosphorylation of PLCγ in the reproductive tract (Fig. 4). Moreover, RNAi-mediated depletion of PLCγ attenuated the pPLCγ signal (Supp. Fig. 2) and egg development (Fig. 3D). These findings pose two immediate questions; how might 20E induce the activation of PLCγ, and how could this PLCγ signal impact egg development? Taking the mechanism of activation first, while classical steroid hormone signaling occurs through the regulation of target gene expression by ligand-activated transcription factors, non-genomic signaling (effects independent of gene expression), often through G protein-coupled plasma membrane receptors, have also been reported in both mammals (27) and insects (28). Our data demonstrate at least a partial role for the conventional EcR (Fig, 4B, D), but we cannot rule out a role for a novel, G protein-coupled receptor for ecdysone in *An. gambiae*.

As discussed below, our data implicate Src family tyrosine kinases in the 20E-dependent activation of PLCγ. Several mechanisms connecting 20E to this kinase family through the traditional EcR have been described. In *Drosophila* 20E increases reactive oxygen species (ROS) (43), an event known to activate Src family kinases (44). Moreover, classical mammalian steroid receptors, such as those for progesterone or estrogen, have been shown to signal through direct (26) or indirect (45, 46) physical interaction with Src family kinases. Notably, the *An. gambiae* EcR (AGAP029539) contains a proline-rich sequence (PPLSPSP), similar to that in the human progesterone receptor proposed to mediate a direct interaction with Src kinases (26). Additionally, the *An. gambiae* genome contains a possible ortholog (AGAP003842) of human PELP1 (Uniprot: Q8IZL8; proline-, glutamic acid- and leucine-rich protein 1), a putative mediator of estrogen receptor/ Src kinase interactions, highlighting the possibility for similar interactions between 20E- and Src kinase signaling in the mosquito.

How might PLCγ activation contribute to the ability of 20E to promote egg development? PLCγ hydrolyses the plasma membrane phospholipid phosphatidylinositol bisphosphate (PIP_2_) to produce two second messengers, diacylglycerol (DAG) and inositol 1,4,5 trisphosphate (IP_3_) which respectively activate protein kinase C (PKC) and trigger release of Ca^2+^ from intracellular stores (47). In *Drosophila* (48) as well as other insects (49, 50) these PLCγ-dependent signals trigger increased uptake of vitellogenin in developing follicles by receptor-mediated endocytosis. PKC has also been reported to phosphorylate the ecdysone co-receptor USP, boosting its capacity to regulate 20E-dependent transcription (28).

PLCγ is also implicated in border cell migration (51, 52), a key event in mid to late oogenesis in *Drosophila* in which a group of 7-9 follicular epithelial cells detach from the epithelium and migrate towards the oocyte before coming to rest close to the oocyte nucleus where they form the micropyle; the structure through which sperm enter and fertilize the oocyte. Females defective in border cell migration exhibit severely depleted egg production and egg fertility (53, 54). The process of border cell migration is triggered by a pulse of 20E signalling but the cell migration itself is directly dependent on the subsequent activation of PLCγ (55) by two oocyte-derived growth factors, EGF (epidermal growth factor) (56) and PVF (platelet-derived growth factor and vascular endothelial growth factor) (57), signaling through their tyrosine kinase-coupled receptors. These events, which to our knowledge have not been studied in *An. gambiae*, position the tyrosine phosphorylation of PLCγ downstream of a 20E signal required for egg development (58, 59), mirroring the 20E-dependent activation of PLCγ we observe after blood feeding. In the future it will be interesting to address whether and when similar mechanisms of interplay between 20E and PLCγ may be involved in mosquito follicle development.

We also addressed the identity of the kinases that could sit upstream of PLCγ. As discussed above, Src kinases are strongly linked to steroid hormone-dependent signaling (26, 27). Moreover, data from mammals (37) as well as other insects (28) demonstrate a specific role for Src family kinases in the phosphorylation of PLCγ. Consistent with this idea, we found that the tyrosine phosphorylation of PLCγ after blood feeding, as well as the subsequent development of eggs, was inhibited by the Src-specific inhibitor SU6656. Identification of the Src kinase family member/s involved in this effect awaits further study.

While the mechanistic details of the signaling relay uncovered here remain to be clarified, the data presented highlight an unexpected connection between the ecdysteroid hormone 20E and the Src-family kinase-dependent tyrosine phosphorylation of PLCγ. Moreover, by establishing a link between tyrosine phosphorylation and egg development our data highlight the utility of targeting tyrosine kinases and tyrosine kinase substrates as a novel strategy to limit reproductive output, and thus vector capacity, in one of the most important vectors of malaria.

## MATERIALS AND METHODS

### Mosquito Rearing

*Anopheles gambiae* mosquitoes of the G3 line (MR4) were maintained under standard MR4 insectary conditions (26–28°C, 65–80% relative humidity, 12:12 hours light/dark photoperiod). Larvae were fed with powdered fish food and carp fish pellets (Tetramin, Tetra Co., Melle, Germany) until pupation. Pupae for colony maintenance were collected in small dishes, and transferred to holding cages (BUGDORM, MegaView Science Co., Taiwan). Experimental pupae were collected and separated by gender by microscopic examination of the terminalia prior to their transfer to cages. Adult mosquitoes were provided with water and 10% w/v glucose *ad libitum*. For blood-fed females, 4-day-old virgin females were fed on human blood (Research Blood Components, Boston, MA, USA) using a Hemotek membrane feeding system.

### RNA-interference

Double-stranded RNA (dsRNA) constructs targeting *PLCγ* (*AGAP029661*), *EcR* (*AGAP029539*), *USP* (*AGAP002095*) were prepared using methods described previously (35). Briefly, PCR primers (see Supplementary Table 1) specific to the gene of interest, designed using E-RNAi webservice (https://www.dkfz.de/signaling/e-rnai3/), were used to generate blunt-ended amplicons from an *An. gambiae* cDNA library which were ligated in to the TOPO 2.1 vector (Thermofisher) and transformed by heat shock in to Top10 competent *E. Coli* (Thermofisher). Plasmid DNA from blue/ white selected, gentamycin-resistant colonies was prepared by midiprep (Thermofisher) and the insert verified by sequencing. Using the purified plasmid as a template, T7 primers (See Supp. Table 1) were used to generate an amplicon of the insert containing T7 RNA polymerase binding sites from which gene-specific dsRNAs were synthesised by *in vitro* transcription (T7 Megascript, Thermofisher). DNase1-digested dsRNAs were purified by phenol/chloroform extraction, washed in ethanol and re-suspended in H_2_O at 10-20μg/μL. A dsRNA targeting *GFP* (not expressed in these mosquitoes) was prepared from an *EGFP*-containing plasmid (60), and used as a negative control.

### dsRNA injection

Sexually mature females (3 or 4 days old) were anaesthetized (Inject+matic sleeper TAS, Geneva, Switzerland) and injected (Nanoject II, Drummond Scientific/Olinto Martelli Srl, Florence, Italy) with 0.69μg (2x 69nL x 5μg/μL) of the appropriate dsRNA then seventy-two hours later females were allowed to take a blood meal. At various points after the blood meal, reproductive tracts (spermatheca, atrium and ovaries) were dissected and prepared for western blot analysis (see below) or were allowed time for eggs to develop completely (48 hours) at which point ovaries were dissected and eggs counted under the microscope.

### Injection of inhibitors, enzymes and 20E

Two hours after blood feeding, females were anaesthetized and injected (2×69nl) with tyrosine kinase inhibitors (all 200μM) SU6656 (CAS 330161-87-0, Cayman Chemical, Ann Arbor, USA), genistein (CAS 446-72-0) or its inactive analog, genistin (CAS 529-59-9, both MilliporeSigma, Burlington, USA), the fungal enzyme ecdysone 22 oxidase (E22O (34); 5μg/ml in PBS) or appropriate vehicle controls; 5% DMSO in PBS (genistein/ genistin/ SU6656) or 5μg/ml BSA in PBS (E22O). His-tagged E22O was cloned and purified from transfected HEK293E cells using nickel affinity chromatography as described in detail elsewhere (2). At various points after blood feeding reproductive tracts (spermatheca, atrium and ovaries) were dissected for Western blot analysis (see below) or allowed time for eggs to develop completely (48h) and eggs counted under the microscope. For injection of 20E (Sigma/ Merck Life Science Srl, Milan, Italy), anaesthetized 3-4-day-old virgin females were injected with 2.45μg 20E (2x 69nL x 37mM) or the same volume of vehicle control (10% EtOH, 5% DMSO in H_2_O) then replaced in the insectary to recover. Heads and reproductive tracts were dissected at various points post injection and analyzed by western blot (see below) for the presence of pPLCγ (reproductive tract) or pJNK (head).

### Western blotting

Samples were prepared and analyzed as previously described in detail (36) with minor modifications. Reproductive tracts (comprising ovaries, atrium and spermatheca), and on occasion, heads and the rest of the body (RoB, remaining carcass minus the midgut which was removed to avoid contamination with ingested blood), were recovered from blood-fed or sugar-fed females (10-15 per point) and placed in 25μl of protein extraction buffer, similar to that previously described (36) but containing a ten-fold higher concentration of SDS (1%). Samples were homogenized, denatured and separated over 4-12% gradient Bis-Tris gels (Thermofisher), then transferred to nitrocellulose membranes (Thermofisher). Membranes were blocked (1h, RT) as described elsewhere (36) using 5% BSA in PBS-Tween or in Odyssey Blocking Buffer (LI-COR Biosciences, Lincoln, NE, USA). Blocked membranes were incubated (1 hour, 1/5000 dilution, RT) with mouse anti-phosphotyrosine (clone 4G10) directly conjugated to HRP (4G10-HRP; Merck/ Millipore, RRID: AB_310779), or unlabelled 4G10^®^ Platinum (Merck/ Millipore, RRID: AB_916371), with rabbit anti-pPLCγ (Cell Signaling Inc, Leiden, Netherlands, RRID: AB_330855) or anti-pJNK (overnight, 4°C, 1/1000 dilution; Cell Signaling Inc, RRID: AB_823588) then washed (4×15 min PBS-Tween). Anti-rabbit or anti-mouse HRP-conjugated secondary antibodies were added (1h, RT, 1/5000 dilution in PBS-Tween containing 5% non-fat milk) then membranes washed again (4×15min, PBS-Tween) and developed using enhanced chemiluminescence (ECL) reagents (Ammersham, Cambridge, UK) and visualized using a Fusion FX chemiluminscence detector (Vilber-Lourmat, Marne-la-Vallée, France) or, on occasion, using the iBright CL1500 imaging system (Thermofisher). Alternatively, the primary antibodies were detected using IRDye 800CW-conjugated Donkey anti-rabbit (RRID: AB_621848) or anti-mouse (RRID: AB_621847) IgG (Li-COR) secondary antibodies and developed using an Odyssey CLx scanner (Li-COR). Membranes were then stripped (ReStore; Thermofisher) and re-probed (1h, RT, 1/10000 diluted in PBS-Tween with 5% non-fat milk) with rat anti-β actin (AbCam, Cambridge, UK, RRID: AB_867488) washed (as above) and incubated with anti-rat-HRP (AbCam, RRID: AB_10680316) or IRDye 680RD-conjugated anti-rat (Li-COR, RRID: AB_10956590) (1h, RT, 1/5000 dilution in PBS-Tween containing 5% non-fat milk) then washed and developed with ECL or using the Odyssey System, as above. Optical density of bands was measured using Image J software (61), normalizing above-background values for pPLCγ, pY or pJNK for the actin signal in each lane. For anti-phosphotyrosine blots the whole lane, rather than a single band, was analyzed. This ratio (pPLCγ/actin, total pY/ actin or pJNK/actin) was used as a measure of ‘relative phosphorylation’ for each sample.

In experiments to address the impact of attenuated 20E or Src kinase signaling on pPLCγ levels, 3-day-old females were injected with ds*EcR* (72 hours before blood feeding), E22O or the Src kinase inhibitor SU6656 (both 2 hours after blood feeding) or with appropriate controls (respectively ds*GFP*, BSA or vehicle) as described above. pPLCγ levels were measured at various points after blood feeding.

### Gene expression analysis by qRT-PCR

Knock down of targeted genes was assessed in whole, unfed mosquitoes, sacrificed just before blood feeding, using qRT-PCR methods described previously (36, 62). Briefly, RNA in homogenates of dissected tissues was purified using Direct-zol RNA miniprep columns (Zymo Research/ Euroclone, Milan, Italy) according to the manufacturer’s instructions. A portion (0.5-1μg) of this material was reverse transcribed as described in detail elsewhere (35). Target gene expression was measured in triplicate 5μL aliquots using Fast SybrGreen Master Mix (Thermofisher), the forward and reverse primers listed in Supplementary Table 2 and a StepOnePlus qRT-PCR thermocycler (Thermofisher). Expression of genes of interest was quantified using the delta CT method using Ribosomal protein L19 (RPL19; AGAP004422), whose expression is not affected by blood feeding (63), as a reference gene. Depletion of target genes was measured using delta delta CT data from *dstarget* vs *dsGFP* controls.

### Immunofluorescence analysis

Dissected ovaries were fixed in 4% paraformaldehyde (30 min, RT), washed (3 × 15 min in PBS) then treated with 3% hydrogen peroxide (5 min, RT) to remove autofluorescence, rinsed (3x in PBS) then simultaneously permeabilized and blocked (2h, RT, 0.5% Triton X100, 1% BSA in PBS). Tissues were then stained with p-Tyr-100 (Cell Signaling, RRID: AB_331228) a mouse anti-phosphotyrosine mAb (1 h, RT, dilution 1:1000,) diluted in blocking solution: 0.5% Triton X100, 1% BSA in PBS), rinsed as above and incubated (1 h, RT) with goat anti-mouse Ig Alexa Fluor 488 (Thermofisher, RRID: AB_2534062), diluted 1:1000 in blocking solution. The tissues were rinsed again and mounted in Vectashield mounting media with DAPI (Vector Labs, Burlingame CA, USA, prod. code H-1200-10). Images were acquired on a Zeiss Axio Observer inverted fluorescent microscope. On occasion, individual images of ovaries taken from vehicle or genistein-treated, blood-fed females were analyzed using Zen software (Carl Zeiss) and the median fluorescence intensity for the whole image measured. For these quantitative analyses, conditions of acquisition (laser power, brightness) were maintained constant between groups, exposure values for the GFP channel were 800ms ± 15% for all images in Figure 1C and 632ms ± 10% for all images in Supplementary Figure 1. Original Czi files, containing metadata for all images used, are available through the Mendeley data repository (see Data availability statement).

### Quantification and statistical analysis

All statistical analyses were performed using Prism 8.0 (GraphPad, La Jolla, USA, RRID: SCR_002798). Non-parametric methods were used except in the case that the normality of all groups analyzed passed a Kolmogorov-Smirnov or Shapiro-Wilk test of normality. In all tests, a significance cut-off of p=0.05 was applied. Where two groups were compared Mann-Whitney or unpaired t tests were performed. Where more than two groups were compared, normal data were analyzed using a 1-way ANOVA test with Dunnett’s multiple comparison correction while for non-normal data a Kruskal-Wallis test with Dunn’s multiple comparison was used. For data characterized by two variables a two-way ANOVA test with Dunnett’s multiple comparison correction was used. In some experiments, differences between ‘control’ and ‘treated’ groups were assessed using one sample t-test testing the null hypothesis that the signal in the ‘control’ should be the same (100%) as the ‘treated’ group. In these cases, normal distribution of the data was verified using a Shapiro-Wilk test.

## Supporting information

Ferracchiato et al Supporting Material

## ACKNOWLEDGEMENTS

The authors gratefully acknowledge Emily Lund, Casey Hill, Kate Thornburg, Sean Scott and Katie Westervelt for help with mosquito rearing; Dr. Duo Peng for bioinformatic assistance and Dr. Vicky Ingham and members of the Catteruccia lab for helpful discussions.

This study was sponsored by European Research Council 7th Research Framework Programme (Project ‘Anorep’ Starting Grant 260897), by a Faculty Research Scholar Award by the Howard Hughes Medical Institute and the Bill and Melinda Gates Foundation (Grant ID: OPP1158190), and by the National Institutes of Health (R01 AI124165, R01 AI104956) to F.C. The content is solely the responsibility of the authors and does not necessarily represent the official views of the National Institutes of Health.

## Data availability

All the unprocessed western blots and immunofluorescence microscopy images featuring directly in the figures or used to build the graphs presented, are available with accompanying metadata through Mendeley data with the following web address: https://data.mendeley.com/datasets/xynhp8mspt/draft?a=1711d135-b9f4-4af9-8d84-bb2626a2fa4e

## AUTHOR CONTRIBUTIONS

Conceptualization, M.P. and F.C.; Methodology, M.P., S.F., F.C.; Investigation, S.F. and M.P.; Formal analysis, S.F. and M.P.; Writing-Original Draft, M.P., S.F. and F.C.; Review and editing, M.P. and F.C.; Visualization, M.P., S.F. and F.C.; Funding Acquisition, F.C.; Resources, F.C.; Supervision, F.C and M.P.

