## Supplementary material for "20E-dependent tyrosine phosphorylation of phospholipase C gamma underpins egg development in the malaria vector *Anopheles gambiae*": Ferracchiato et al Supporting Material

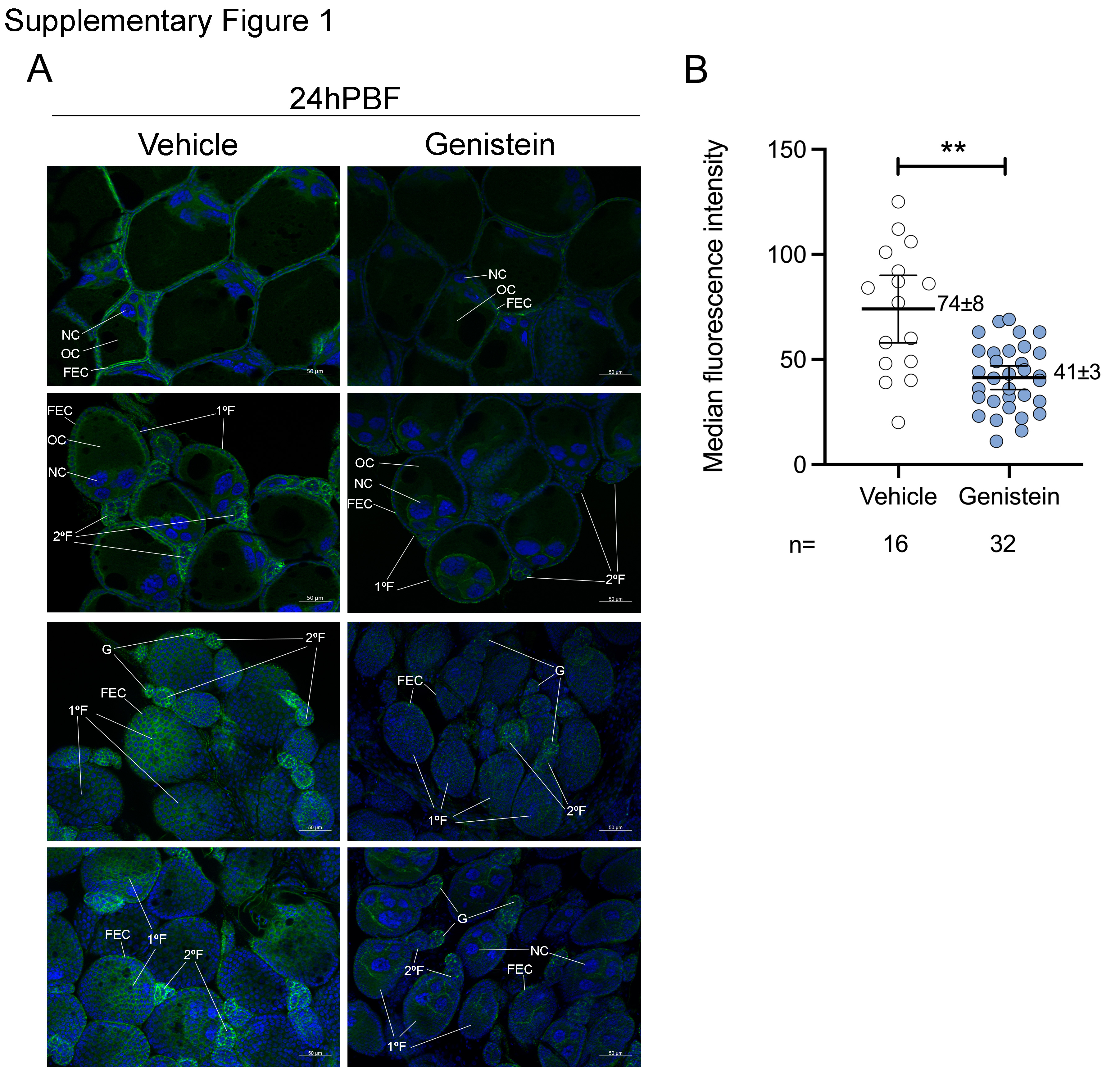

**­­Supplementary Figure 1. Tyrosine phosphorylation at 24 hPBF is inhibited by Genistein injection.** A. Three-day-old females were allowed to take a blood meal and 2 hours later injected with Genistein (200μM) or vehicle control and ovaries dissected 24h PBF. Phosphotyrosine (shown in green) was visualized using an anti-phosphotyrosine primary antibody and an Alexa 488-coupled secondary antibody; cellular nuclei were stained with DAPI (shown in blue). Primary follicles (1ºF), secondary follicles (2ºF) and germaria (G), as well as follicular epithelial cells (FEC), developing oocytes (OC) and nurse cells (NC) are indicated. The images presented were acquired using the same exposure conditions, are from the same group of mosquitoes and are representative of two independent experiments. B. Each point represents the median fluorescence intensity of an individual 20x image. The total number of images analysed is indicated below (n). Images were taken from 5-7 different mosquitoes from each treatment group. Differences between groups were assessed using an unpaired t test. ** denotes­ p<0.01. The data are representative of two independent experiments.

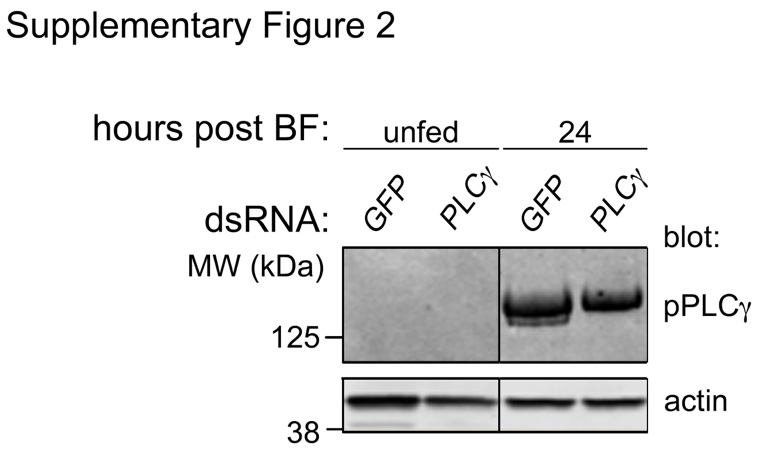

**Supplementary Figure 2. Blood feeding induces tyrosine phosphorylation of PLCγ.** Three-day-old females were injected with ds*PLCγ* or ds*GFP* then 72 hours later maintained on sugar alone (unfed) or allowed to take a blood meal. Twenty-four hours after blood feeding, reproductive tracts (10/point) were dissected from all groups and analyzed in parallel for phospho PLC gamma (pPLCγ) and actin by cutting the membrane above the 50kDa marker. A single representative western blot is shown of the three *dsPLCγ* depletion experiments presented in Figures 3C and 3D. In the original image the lane order of ds*GFP* and ds*PLCγ* treatments was reversed. The line separating the unfed and 24 hPBF time points indicates the digital alteration of the original lane order to maintain consistency with other Figures.

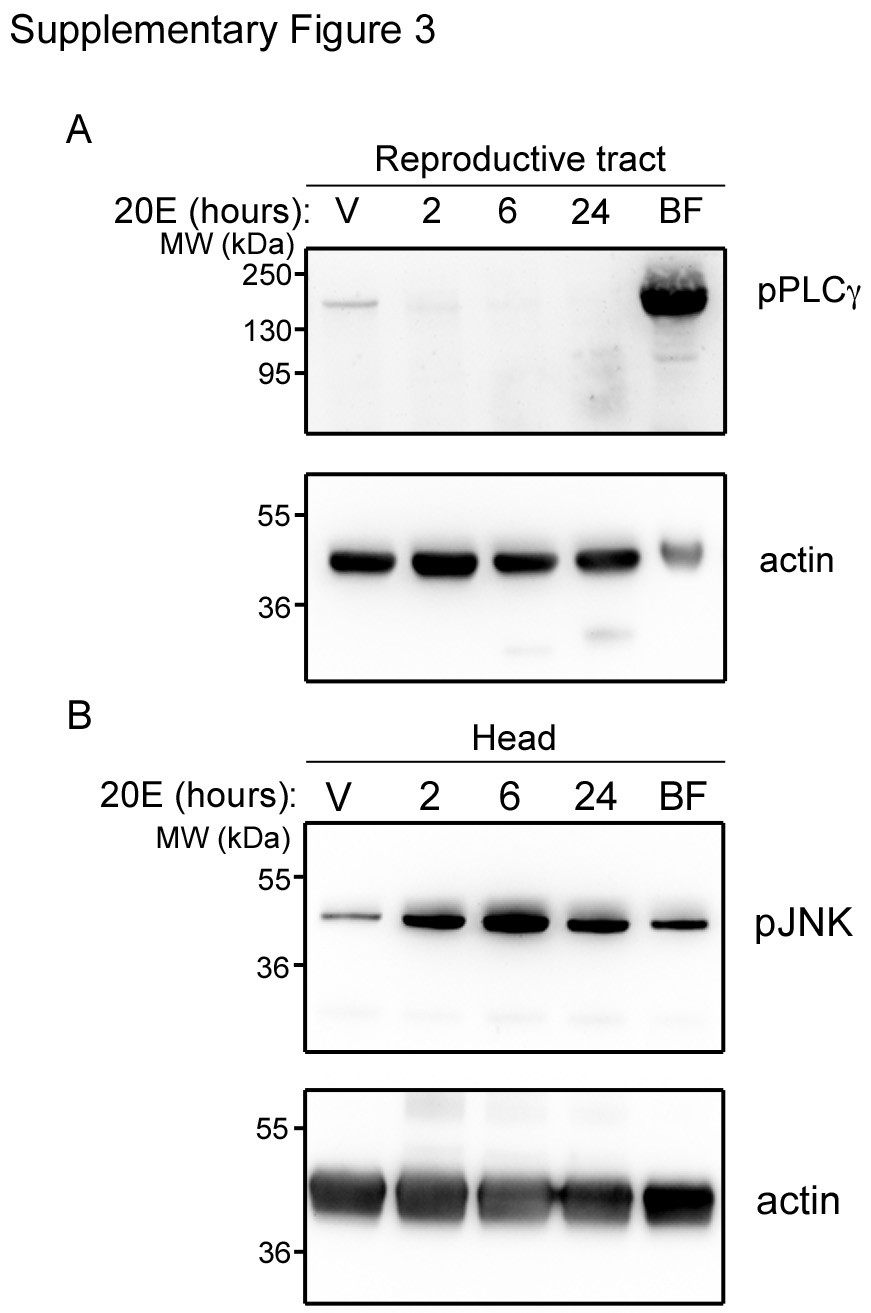

**Supplementary Figure 3. Injected 20E does not induce pPLCγ.** A. Reproductive tracts (comprising atrium, spermatheca and ovaries), or heads (B) were dissected from unfed females injected with 20 hydroxyecdysone (20E, 37mM) at the time points indicated, at 24 hours post injection of a vehicle-injected control (V) or 24 hours after blood feeding (BF). Pooled tissues (10/ time point), were analyzed by western blot for phospho-PLCγ (pPLCγ) or pJNK as indicated, then stripped and reprobed for actin. A representative blot of two similar experiments is shown.

**Supplementary Table 1**. Primer sequences used to generate dsRNAs.

| **Target** | **Primer name** | **Sequence (5’→3’)** |
| --- | --- | --- |
| PLCγ  AGAP029661 | PLCγ F  PLCγ R | ACCAGCGGCTGATACTGC  GCTCATGTCCTGGTACACCG |
| EcR  AGAP029539 | EcR F  EcR R | CTGCTCCAGTGAGGTGATGA  GGCAGCTTACGGTTCTTCAG |
| USP  AGAP002095 | Usp F  Usp R | AGAAGGAGAAACCGATGCTG  AAATGTCCGGCTTCAGGTC |
| T7 | T7 Fwd | aatacgactcactatagggCCGCCAGTG  TGCTGGAA |
| T7 | T7 Rev | taatacgactcactatagggCCAGTGTGAT  GGATATCTGCAGAA |

**Supplementary Table 2**. Primer sequences used for RT-qPCR analysis.

| **Target** | **Primer name** | **Sequence (5’→3’)** |
| --- | --- | --- |
| RPL19 AGAP004422 | RPL19 F  RPL19 R | CCAACTCGCGACAAAACATTC  ACCGGCTTCTTGATGATCAGA |
| PLCγ  AGAP029661 | PLCγ F  PLCγ R | CACTTTTTCGTGCTGACGCA  TCTCTTCGCGCTCATCATCC |
| GFP | GFP F  GFP R | TGTTCTGCTGGTAGTGGTCG  ACGTAAACGGCCACAAGTTC |
| EcR  AGAP029539 | EcR F  EcR R | CTGCTCCAGTGAGGTGATGA  GGCAGCTTACGGTTCTTCAG |
| USP  AGAP002095 | Usp F  Usp R | GGAAGCAATGGAGGTGGAG  CATAGAATTCTGGCCAACGC |

All qRT-PCR primers were used at 300nM except RPL19 Rev which was used at 900nM.

**FULL LENGTH WESTERN BLOTS**

Figure 1A

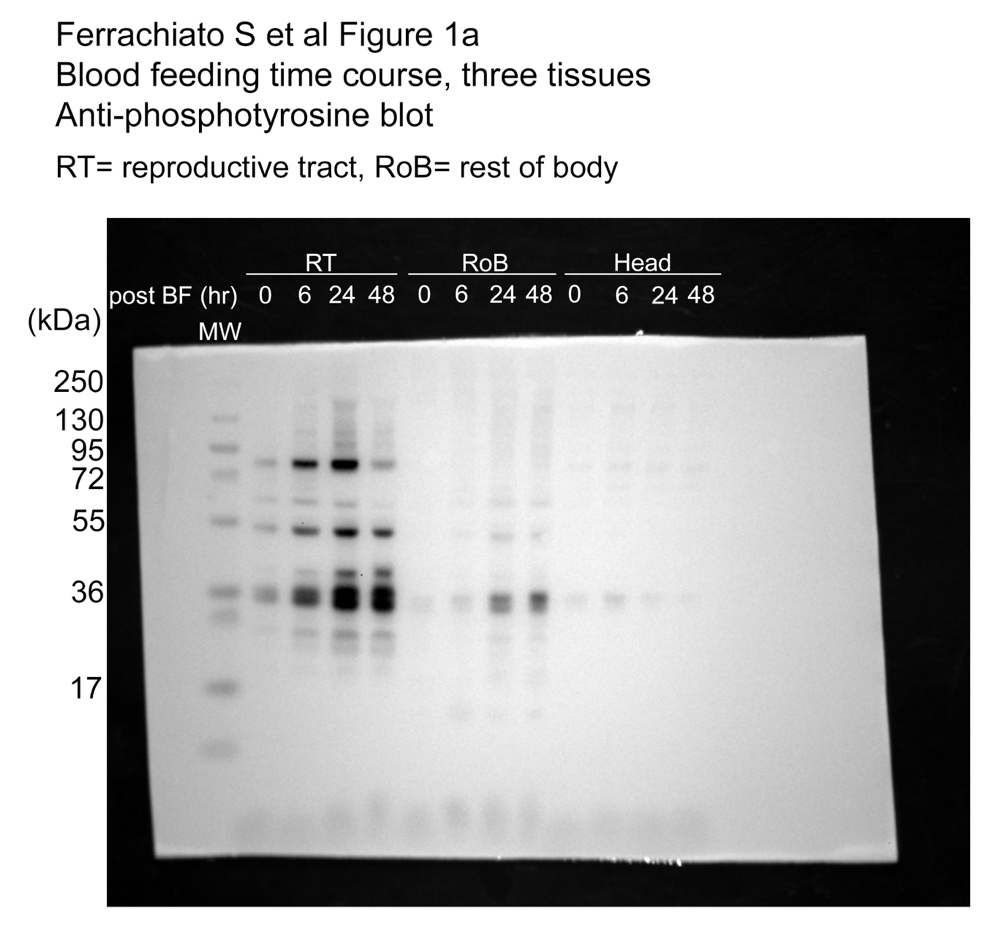

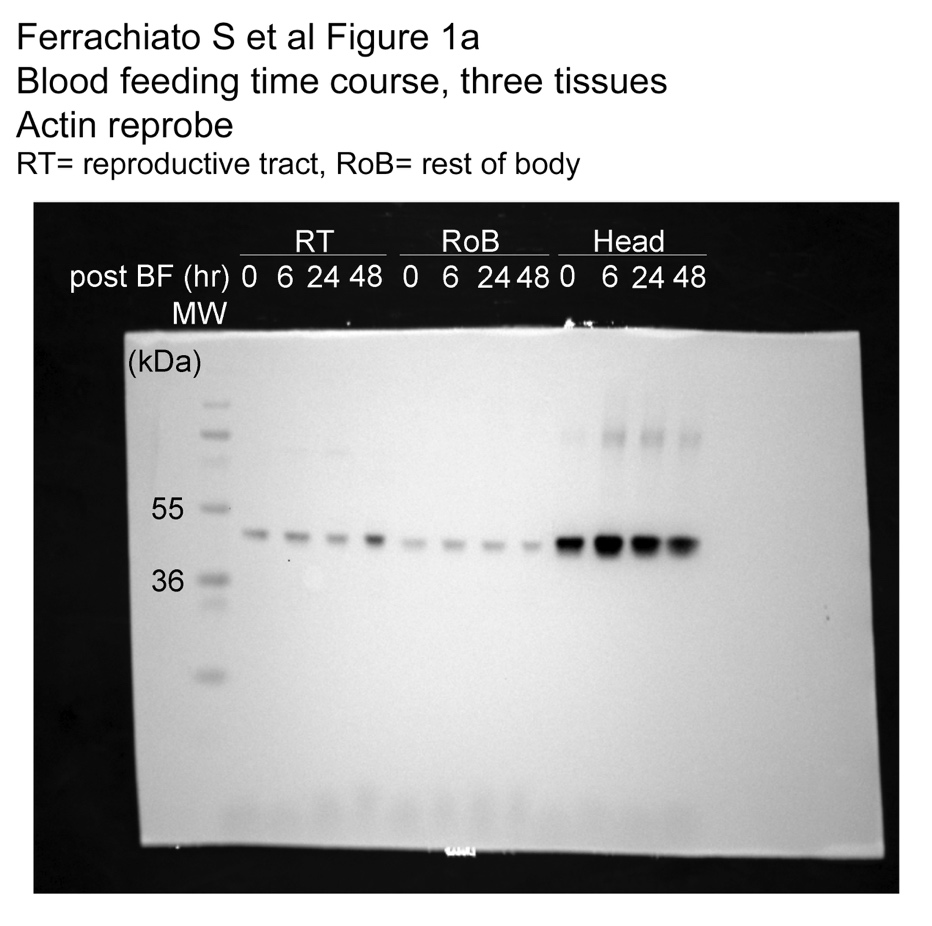

Figure 2A

NB Genistein 48, 24 h (labelled in grey) presented in the opposite orientation in the final Figure.

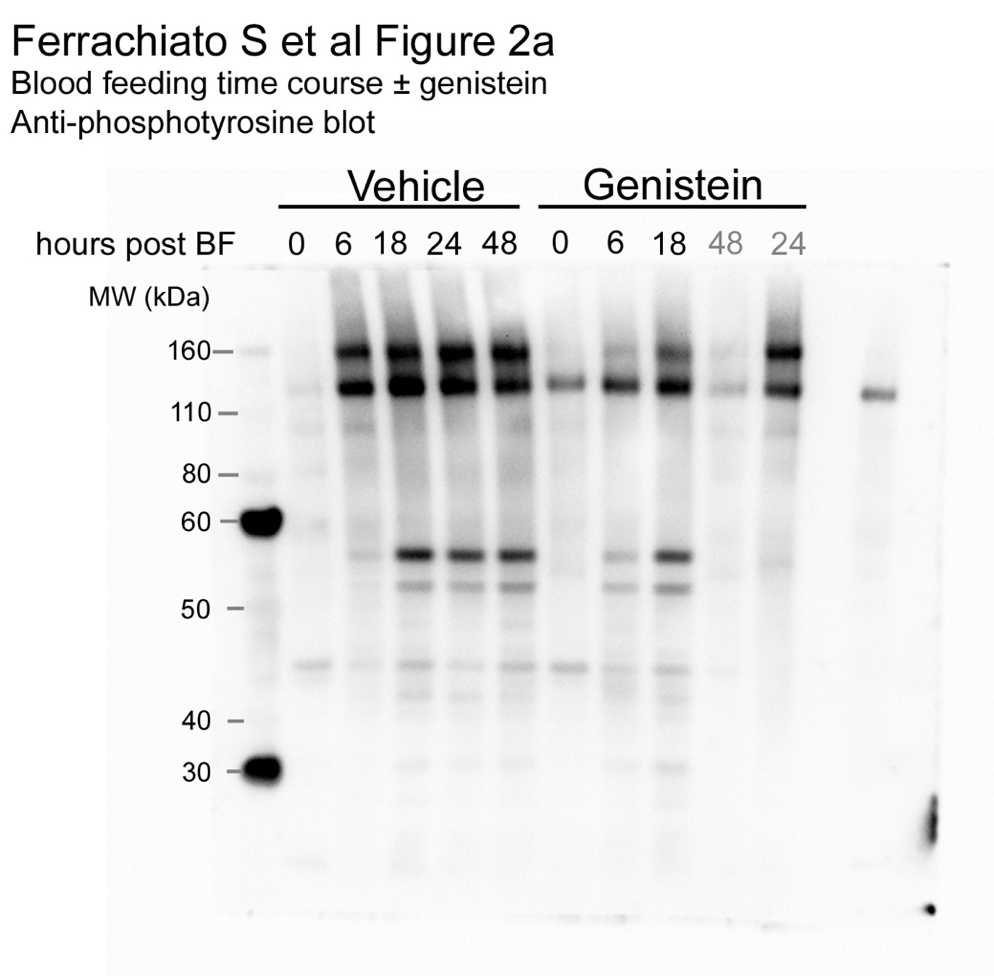

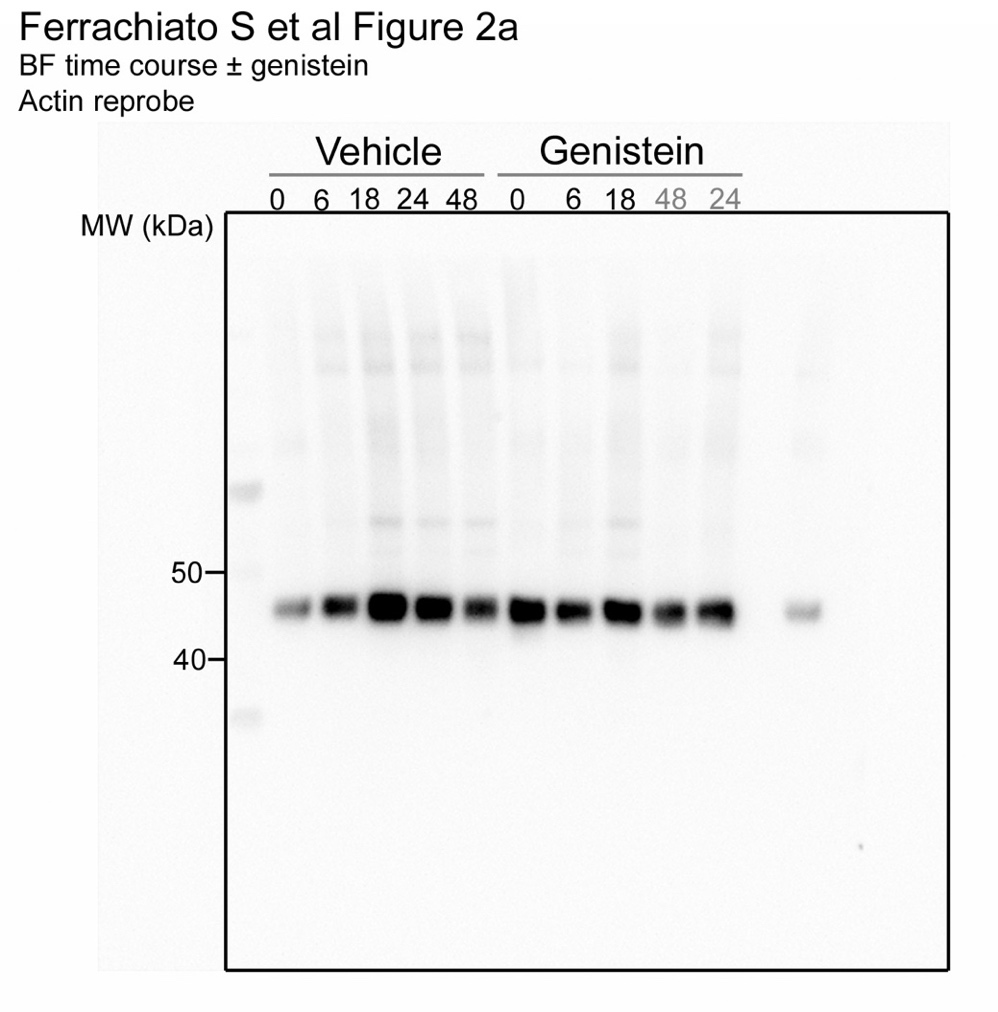

Figure 3A

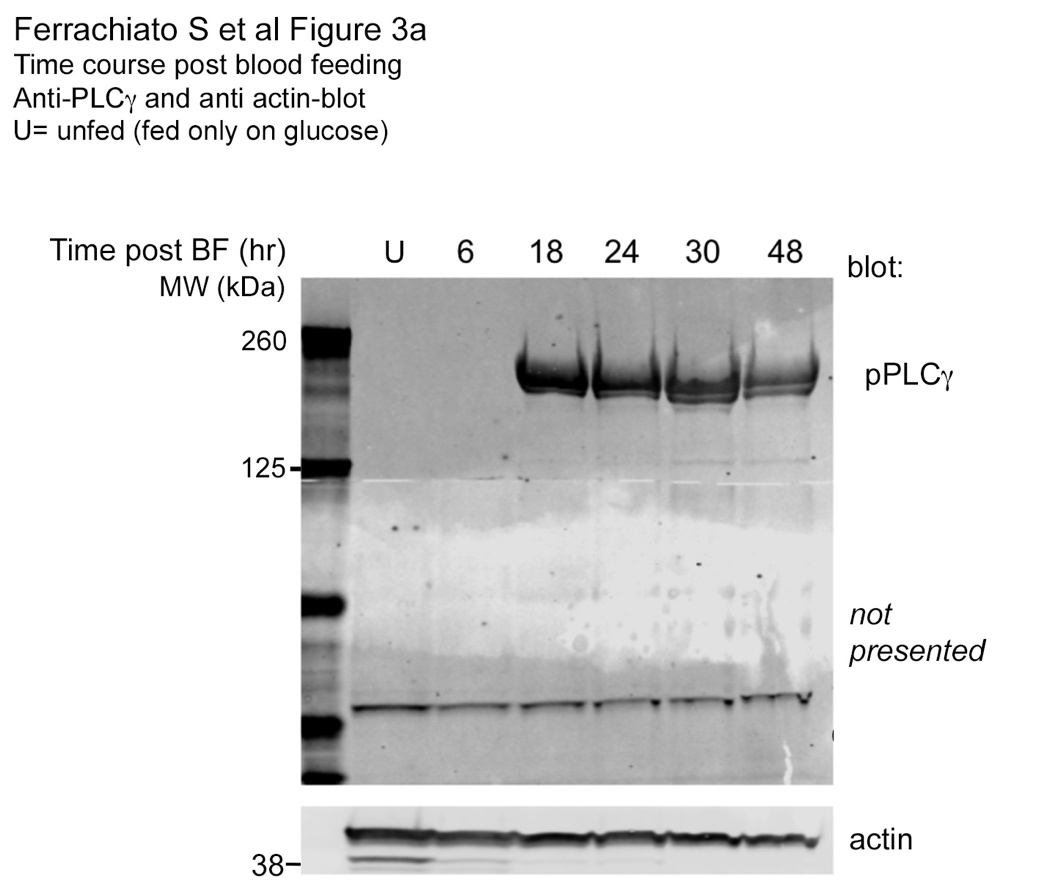

Figure 4A

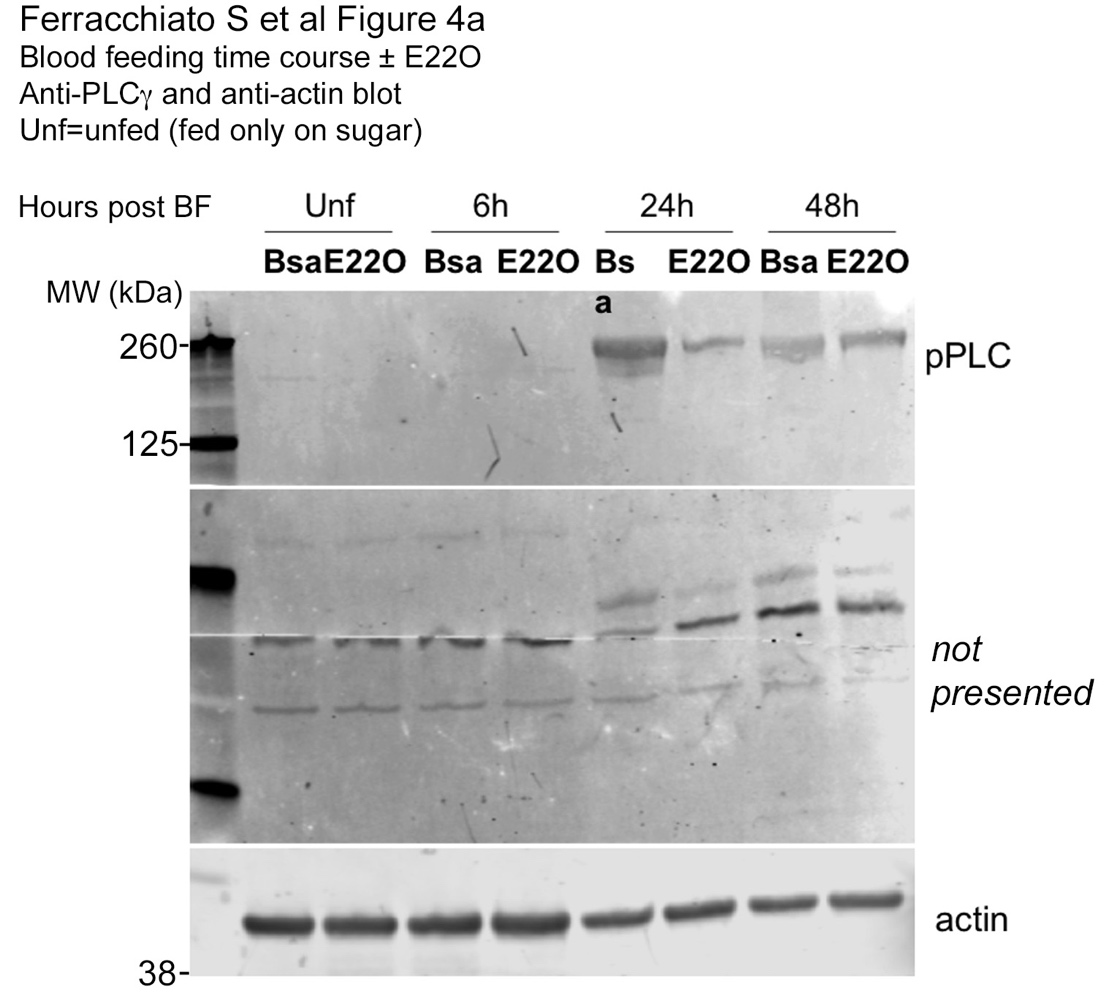

Figure 4B

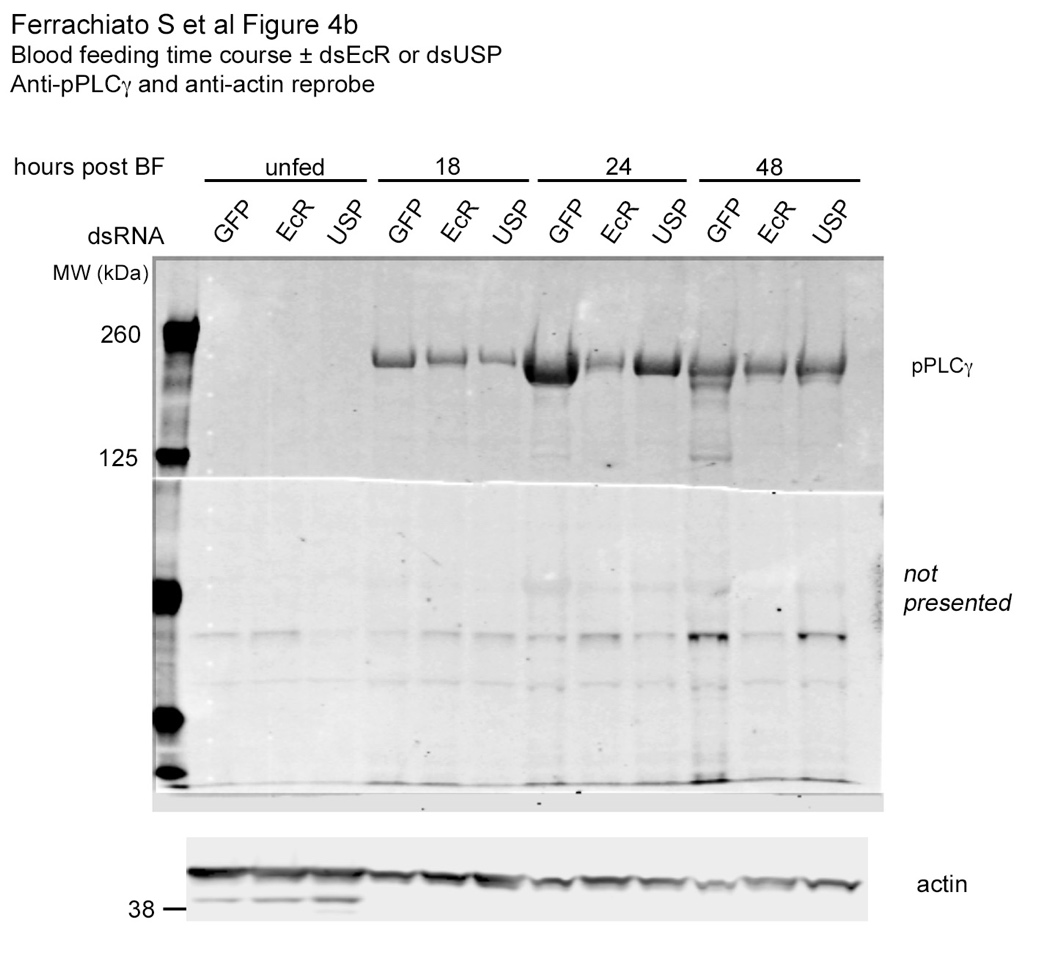

Figure 5A

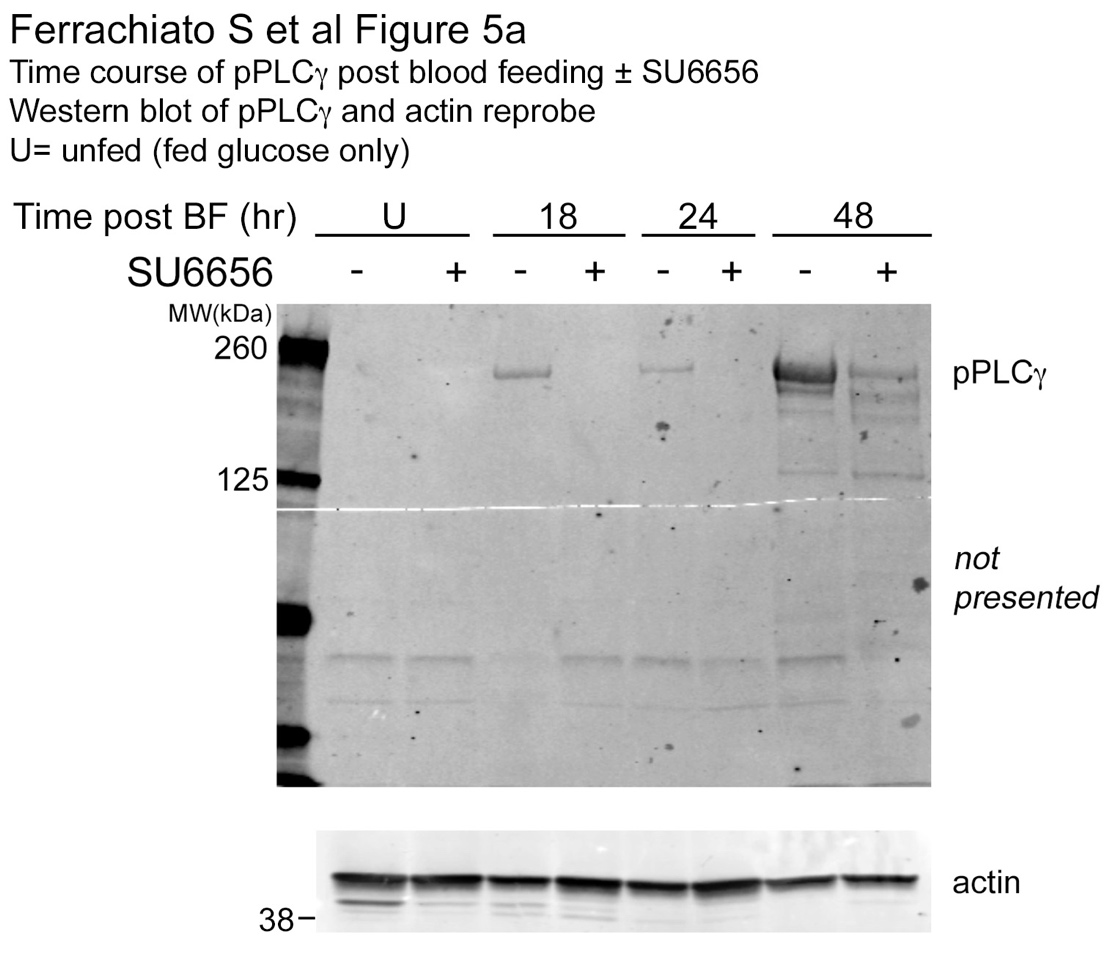

Supplementary Figure 2

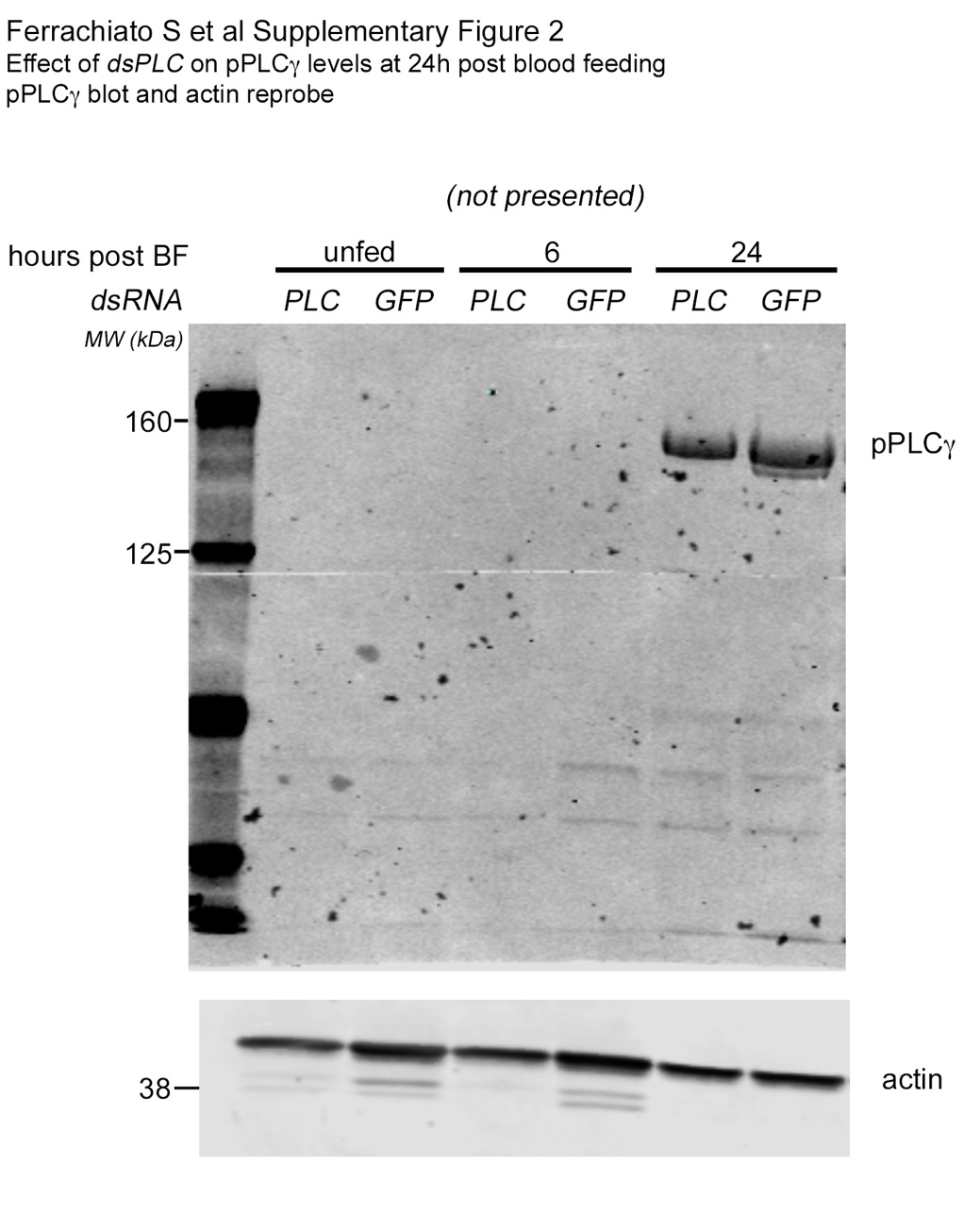

Supplementary Figure 3A (Reproductive Tract)

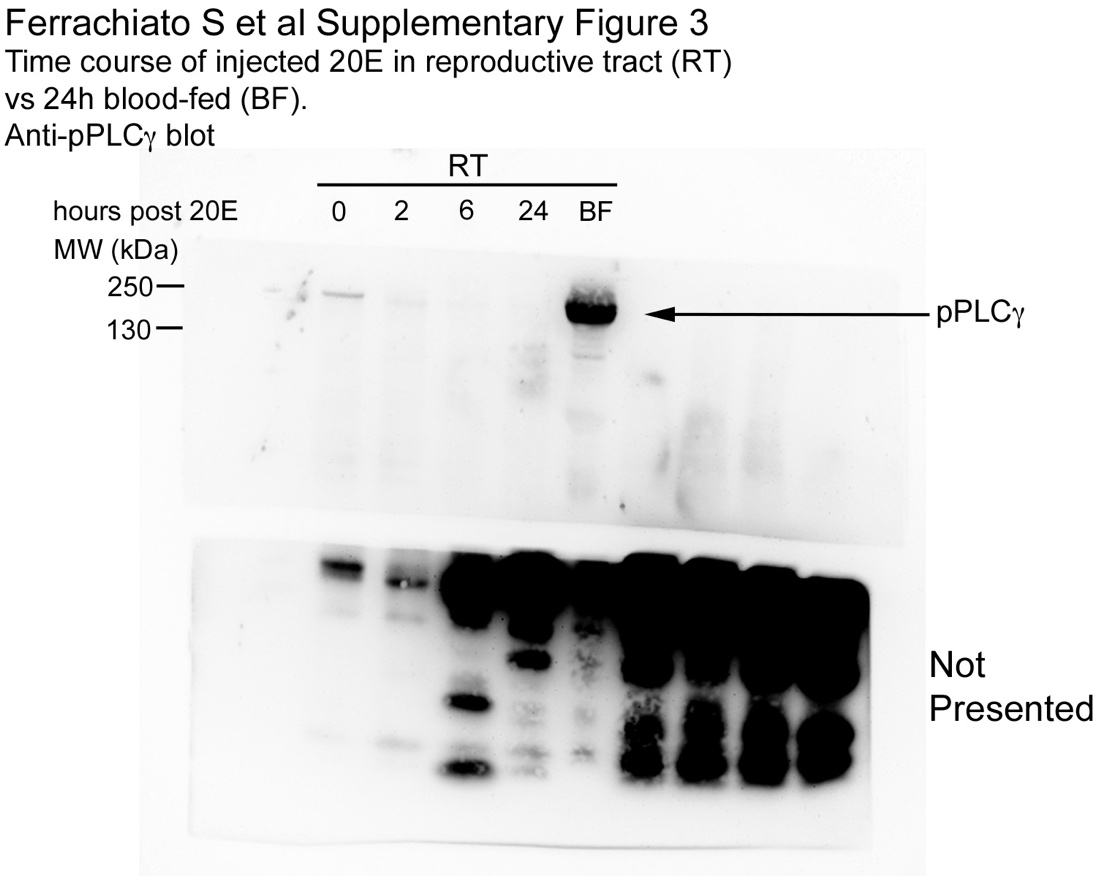

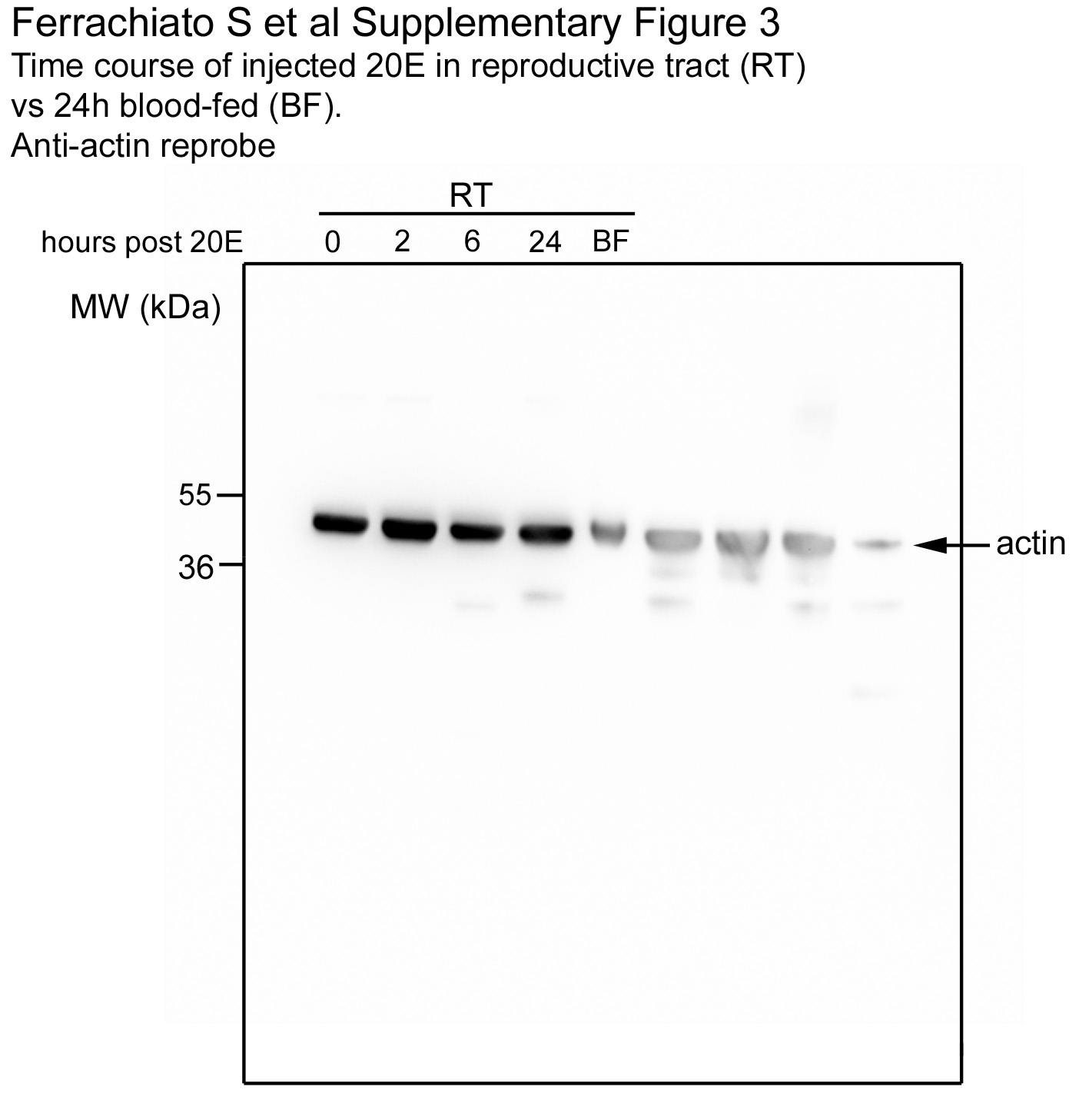

Supplementary Figure 3B (Head)

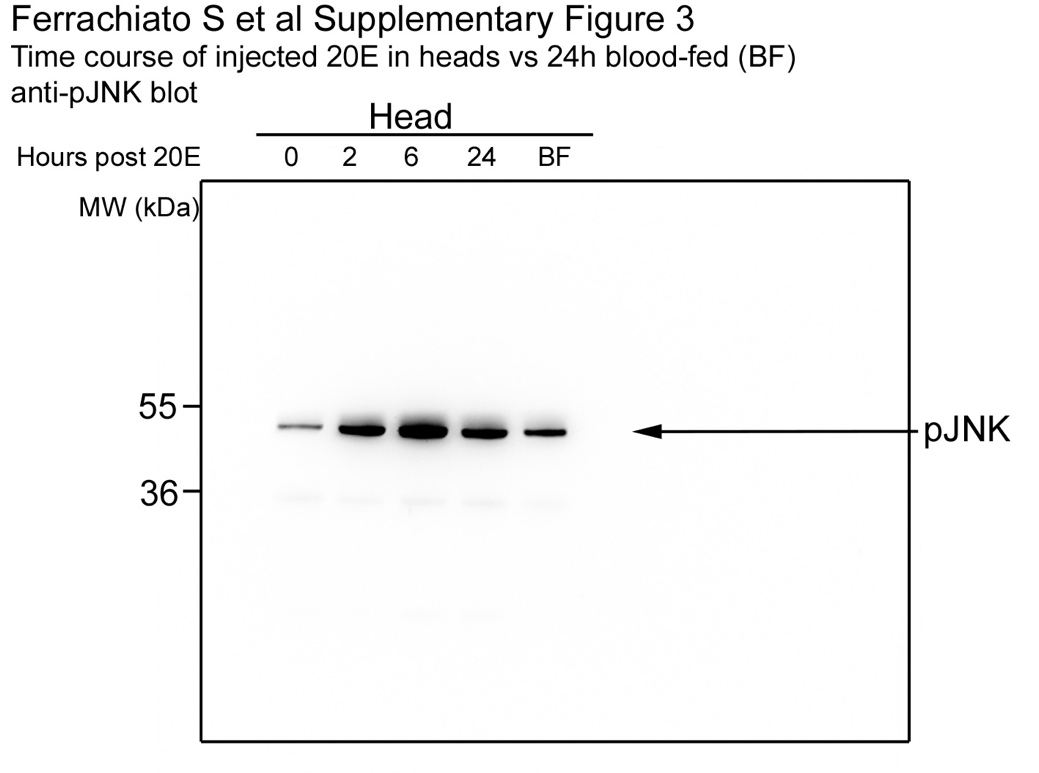

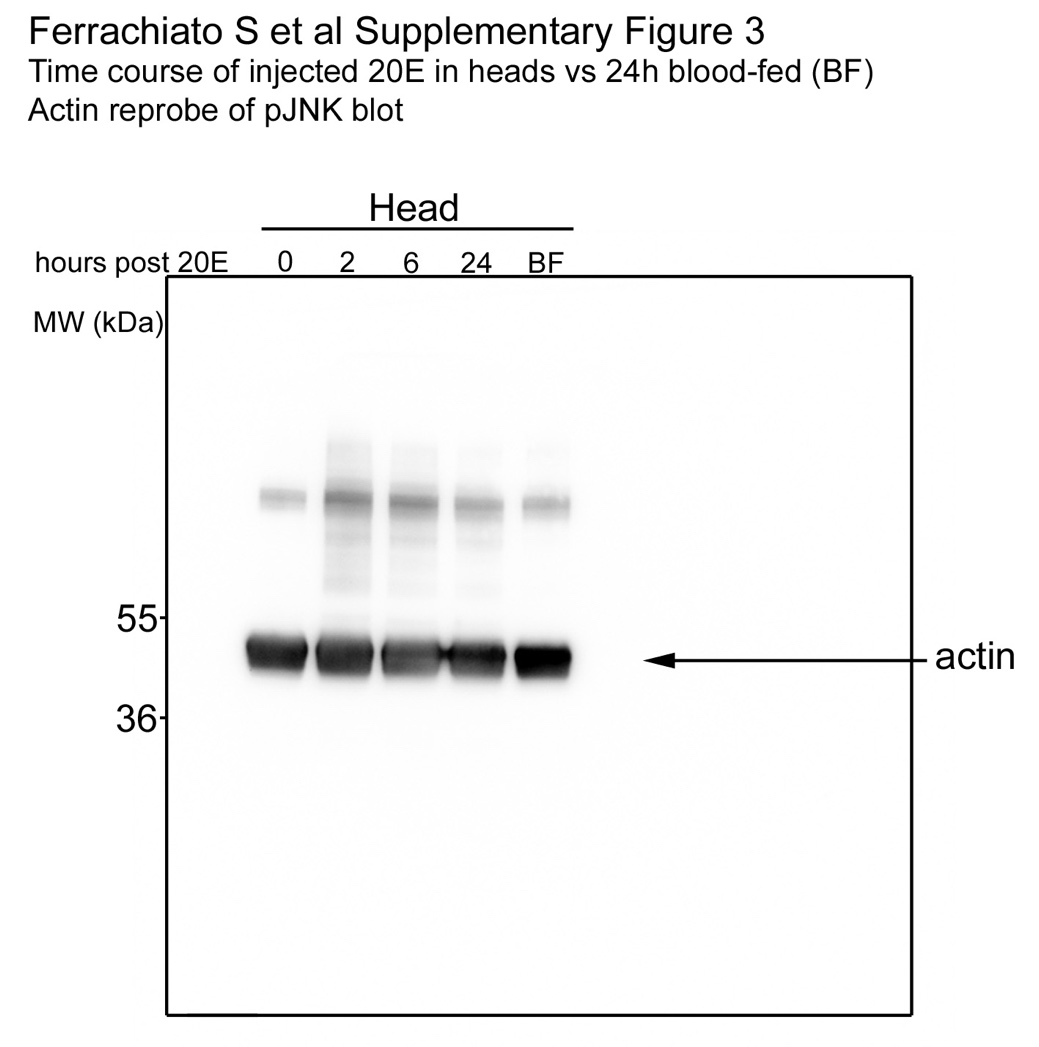
